# The nucleotide binding domain of NRC-dependent disease resistance proteins is sufficient to activate downstream helper NLR oligomerization and immune signaling

**DOI:** 10.1101/2023.11.30.569466

**Authors:** Mauricio P. Contreras, Hsuan Pai, Rebecca Thompson, Jules Claeys, Hiroaki Adachi, Sophien Kamoun

## Abstract

Nucleotide-binding domain and leucine-rich repeat (NLR) proteins with pathogen sensor activities have evolved to initiate immune signaling by activating helper NLRs. However, the mechanisms underpinning helper NLR activation by sensor NLRs remain poorly understood. Although coiled-coil (CC) type sensor NLRs such as the *Potato virus X* disease resistance protein Rx have been shown to activate the oligomerization of their downstream helpers NRC2 and NRC4, the domains involved in sensor-helper signaling are not known. Here, we show that the nucleotide binding (NB) domain within the NB-ARC of the *Potato virus X* disease resistance protein Rx is necessary and sufficient for oligomerization and immune signaling of downstream helper NLRs. In addition, the NB domains of the disease resistance proteins Gpa2 (cyst nematode resistance), Rpi-amr1, Rpi-amr3 (oomycete resistance) and Sw-5b (virus resistance) are also sufficient to activate their respective downstream NRC helpers. Moreover, the NB domain of Rx and its helper NRC2 form a minimal functional unit that can be transferred from solanaceous plants (lamiids) to the Campanulid species lettuce (*Lactuca sativa*). Our results challenge the prevailing paradigm that NLR proteins exclusively signal via their N-terminal domains and reveal a signaling activity for the NB domain of NRC-dependent sensor NLRs. We propose a model in which helper NLRs monitor the status of the NB domain of their upstream sensors.

## Introduction

NLRs (nucleotide binding and leucine-rich repeat) are intracellular innate immune receptors of eukaryotes and prokaryotes (Chou *et al*, 2023; Contreras *et al*, 2023a). In plants, they directly or indirectly sense pathogen virulence proteins, termed effectors, and mediate robust immune signaling and disease resistance (Contreras *et al*., 2023a; Duxbury *et al*, 2021). Plant NLRs exhibit a conserved domain architecture, consisting of an N-terminal domain, a central NB-ARC (nucleotide-binding domain shared by APAF1, R gene product and CED-4) and a C-terminal leucine-rich repeat (LRR) region (Kourelis *et al*, 2021). The NB-ARC can be subdivided into a nucleotide-binding (NB) domain, a helical domain 1 (HD1) and a winged-helix domain (WHD) (Chou *et al*., 2023; Förderer & Kourelis, 2023). Based on their N-terminal domain features, angiosperm NLRs can be broadly categorized into Toll/Interleukin-1 receptor (TIR)-type, coiled coil (CC)-type, G10 subclade coiled coil (CC_G10_)-type and RPW8 coiled coil (CC_R_)-type, which follow the NB-ARC-based NLR phylogeny (Kourelis *et al*., 2021). Some NLRs can function as individual units, termed singletons, mediating both pathogen perception and immune signaling (Adachi *et al*, 2019b). However, coevolution with pathogens has led to functionally specialized NLRs, where pathogen perception and immune signaling become uncoupled. In these cases, one NLR acts as a pathogen sensor and relies on a downstream helper NLR to mediate immune signaling and disease resistance. Sensors and helpers can function as genetically linked pairs or in higher order configurations that can include genetically unlinked receptor networks (Adachi & Kamoun, 2022; Wu *et al*, 2018). However, in contrast to singleton NLRs (Contreras *et al*., 2023a), the activation mechanisms of paired and networked NLR are less understood. In particular, the molecular mechanisms underpinning CC-NLR sensor-helper communication are not known.

The current paradigm for NLR activation is that effector perception leads to oligomerization and induced-proximity of the N-terminal domains (Contreras *et al*., 2023a; Duxbury *et al*., 2021). For CC-NLRs such as Arabidopsis ZAR1 and wheat Sr35, effector recognition triggers conformational changes in inactive NLR monomers which lead to the assembly of a pentameric resistosome complex (Förderer *et al*, 2022; Wang *et al*, 2019; Zhao *et al*, 2022). In the resistosome, the N-terminal α1-helices of the CC domains come together to form a funnel-like structure that inserts into the plasma membrane, presumably to perturb membrane integrity and act as a calcium channel (Bi *et al*, 2021; Wang *et al*., 2019). The α1-helices of ZAR1, Sr35 and around 20% of all angiosperm CC-NLRs are defined by the MADA motif, which is crucial for cell death induction (Adachi *et al*, 2019a). In some cases, the CC domain or the α1-helix on their own are sufficient to initiate cell death induction, presumably via the assembly of a resistosome-like structure that retains the capacity to de-stabilize the plasma membrane and mediate calcium influx (Adachi *et al*., 2019a; Bentham *et al*, 2018; Bi *et al*., 2021; Förderer *et al*., 2022). The CC domain is, therefore, considered as the executor domain of downstream signaling in singleton or helper CC-NLRs. In contrast, sensor NLRs have lost this activity throughout evolution due to degeneration of the MADA motif and even acquisition of novel N-terminal domains upstream of the CC domain (Adachi *et al*., 2019b; Contreras *et al*., 2023a). How sensor-helper sub-functionalization shaped the activities of paired and networked CC-NLR domains is not well understood. In particular, the domains of sensor NLRs that mediate signal transduction and communication with downstream helper NLRs are unknown.

In the Solanaceae, the NLRs required for cell death (NRC) network is composed of multiple sensor CC-NLRs and cell-surface receptors which genetically require downstream helper CC-NLRs, known as NRCs (Kourelis *et al*, 2022; Wu *et al*, 2017). This immune receptor network is of great agronomical importance, mediating immunity to diverse plant pathogenic oomycetes, fungi, nematodes, viruses, bacteria, and insects (Derevnina *et al*, 2021; Kourelis *et al*., 2022; Wu *et al*., 2017). All NRC-dependent sensors fall into an expanded phylogenetic clade that includes many well-known disease resistance proteins while the NRC helpers form a tight and well-supported sister clade (Wu *et al*., 2017). The NRC-dependent sensor NLRs themselves cluster into two distinct phylogenetic groups, the Rx-type and the Solanaceous domain (SD)-type clade (Contreras *et al*., 2023a). Interestingly, not all NRC-dependent sensors can activate all helpers, reflecting a degree of specificity that has probably resulted from sensor-effector and sensor-helper co-evolution (Adachi *et al*., 2019b; Contreras *et al*., 2023a). For instance, the disease resistance sensor NLRs Rx, Sw-5b, Gpa2, Bs2 and Rpi-amr3 can signal interchangeably through NRC2, NRC3 or NRC4, whereas Rpi-amr1e signals through NRC2 and NRC3 but not NRC4 and Rpi-blb2 and Mi-1.2 signal through NRC4 but not NRC2 or NRC3 (Ahn *et al*, 2023; Contreras *et al*, 2023c; Derevnina *et al*., 2021; Lin *et al*, 2022; Witek *et al*, 2021; Wu *et al*., 2017). This complex configuration of many-to-one and one-to-many functional sensor–helper connections likely contributes to increased robustness and evolvability of the NRC immune network (Adachi *et al*., 2019b; Contreras *et al*., 2023a; Wu *et al*., 2018).

Phylogenomic analyses revealed that the NRC superclade emerged from a pair of genetically linked NLRs prior to the split between the Asterid and Caryophyllales lineages over 100 million years ago (mya) (Wu *et al*., 2017). Two recent studies on NRC network diversity and macroevolution across Asterid plants mapped out massive expansions of family-specific NRC sensors and helpers in the Lamiid lineage within asterids, which include the Solanacaeae, in contrast to the more broadly conserved NRC0 helpers and their genetically linked sensors (Goh *et al*, 2023; Sakai *et al*, 2023). These studies also highlighted an overall lack of sensor-helper interchangeability across taxa (Goh *et al*., 2023; Sakai *et al*., 2023). How sensor-helper compatibility and specificity have evolved is not understood, given that the molecular determinants for sensor-helper specificity are not known.

In previous studies, we proposed an activation-and-release biochemical model for sensor-helper activation in the NRC network: Pathogen-activation of NRC-dependent sensor NLRs leads to homo-oligomerization of their NRC helpers into resistosome complexes which accumulate at the plasma membrane, separate from the sensors that activated them (Contreras *et al*., 2023c). Specifically, the sensor NLR proteins Rx (virus resistance), Bs2 (bacterial resistance) and Rpi-amr3 (oomycete resistance) can trigger oligomerization of NRC2 and NRC4 (Ahn *et al*., 2023; Contreras *et al*, 2023b; Contreras *et al*., 2023c). In contrast, Rpi-amr1e (oomycete resistance) can oligomerize NRC2 but not NRC4 and Rpi-blb2 (oomycete resistance) can oligomerize NRC4 but not NRC2 (Ahn *et al*., 2023; Contreras *et al*., 2023c). In addition, this activation mechanism appears to be conserved in the tomato (*S. lycopersicum*) NRC0 helper and its genetically linked sensor (Sakai *et al*., 2023). This indicates that the activation-and-release mechanism is likely conserved across the NRC superclade. However, the nature of the signal relayed by sensor NLRs to initiate NRC oligomerization is unknown.

The NRC-dependent sensor NLR Rx mediates extreme immunity to *Potato virus X* (PVX) by recognizing its coat protein (CP) via an unknown mechanism (Bendahmane *et al*, 1999; Bendahmane *et al*, 1995; Tameling & Baulcombe, 2007). In a study published over 20 years ago, Moffett and colleagues showed that Rx can perceive PVX CP and cause hypersensitive cell death when expressed *in trans* as two separate halves in the model plant *Nicotiana benthamiana* (Moffett *et al*, 2002). This work resulted in a a highly influential mechanistic model, which postulated that multiple intramolecular interactions maintain Rx in an inactive state. Upon effector perception, these intramolecular interactions are relieved via conformational changes within the NLR protein, resulting in dissociation of the Rx halves and initiation of immune signaling (Moffett *et al*., 2002). In a follow up study, Rairdan and colleagues went further, demonstrating that a ∼150 amino acid region of Rx corresponding to the NB domain (Rx^NB^) is sufficient to trigger the hypersensitive cell death when expressed in *N. benthamiana* (Rairdan *et al*, 2008). They proposed that upon activation, intramolecular rearrangements in the Rx protein release the NB domain from negative regulation by the CC and LRR domains to enable signaling and cell death induction. However, these findings have remained puzzling considering that the prevailing models of CC-NLR activation assign the singaling activity to the very N-terminal CC and not the NB domain as discussed above. Moreover, the work by Moffett, Rairdan and colleagues predates the discovery of Rx dependence on the NRC2, NRC3 or NRC4 helpers to cause the hypersensitive cell death and confer immunity and resistance to *Potato virus X* (Derevnina *et al*., 2021; Wu *et al*., 2017). Whether the deconstructed and minimal Rx domains signal via the NRC helpers is unknown.

In this study, we revisited the work of Moffett, Rairdan and colleagues on Rx in the context of the NRC immune receptor network. We show that the Rx halves expressed *in trans* as well as Rx^NB^-eGFP can activate downstream helper NLR oligomerization and immune signaling. In addition, the NB domain truncations of the Rx-type sensors Gpa2, Rpi-amr1e and Rpi-amr3 as well as the SD-type sensor Sw-5b can also activate NRC-dependent cell death and exhibit the same downstream helper-specificities of their full-length counterparts. Finally, we show that full-length PVX CP-activated Rx and Rx^NB^-eGFP both trigger cell death when co-expressed with NRC2 in the unrelated Campanulid lettuce, suggesting that Rx^NB^ and NRCs are likely a minimal two-component system that can be transferred across plant taxa. Our findings reveal a novel signaling role for the NB domain in NRC-dependent sensor NLRs, serving as a minimal signal for helper activation. We suggest a model where sensor NLRs undergo conditional NB-ARC domain rearrangements upon effector perception, exposing the NB domain to activate their downstream NRC helpers and triggering oligomerization into resistosomes.

## Results

### The hypersensitive response caused by expression of Rx halves *in trans* with the coat protein (CP) of *Potato virus X* (PVX) is dependent on NRC helpers

To test whether the Rx halves expressed *in trans* activate cell death via NRC-dependent pathways, we expressed Rx^CCNBARC^ and Rx^LRR^ or full-length Rx with or without PVX CP in leaves of WT, *nrc2/3*, *nrc4a/b* or *nrc2/3/4* KO *N. benthamiana* (**Figure 1A**). The constitutively active NbZAR1^D481V^ variant was used as a control for NRC-independent cell death (Harant *et al*, 2022). Like full-length Rx, cell death mediated by Rx^CCNBARC^ and Rx^LRR^ expressed *in trans* was only abolished in the *nrc2/3/4* KO *N. benthamiana* background but not in the *nrc2/3* or *nrc4a/b* KO backgrounds, suggesting that it is also NRC2/3/4-dependent (**Figure 1B**). We also carried out complementation assays in *nrc2/3/4* KO *N. benthamiana* lines to further confirm the NRC-dependency of cell death mediated by Rx^CCNBARC^ and Rx^LRR^. We co-expressed the two Rx halves with PVX CP in the *nrc2/3/4* background and complemented with NbNRC2, NbNRC3 and NbNRC4. We included tomato (*S. lycopersicum*) SlNRC0, an NRC that full length Rx is unable to signal through, as an independent negative control for complementation (Sakai *et al*., 2023). Like full-length Rx, the cell death in response to PVX CP mediated by the Rx^CCNBARC^ and Rx^LRR^ halves expressed *in trans* was restored upon complementation with NbNRC2, NbNRC3 and NbNRC4, but not SlNRC0 (**Figure 1C**).

**Figure 1:**
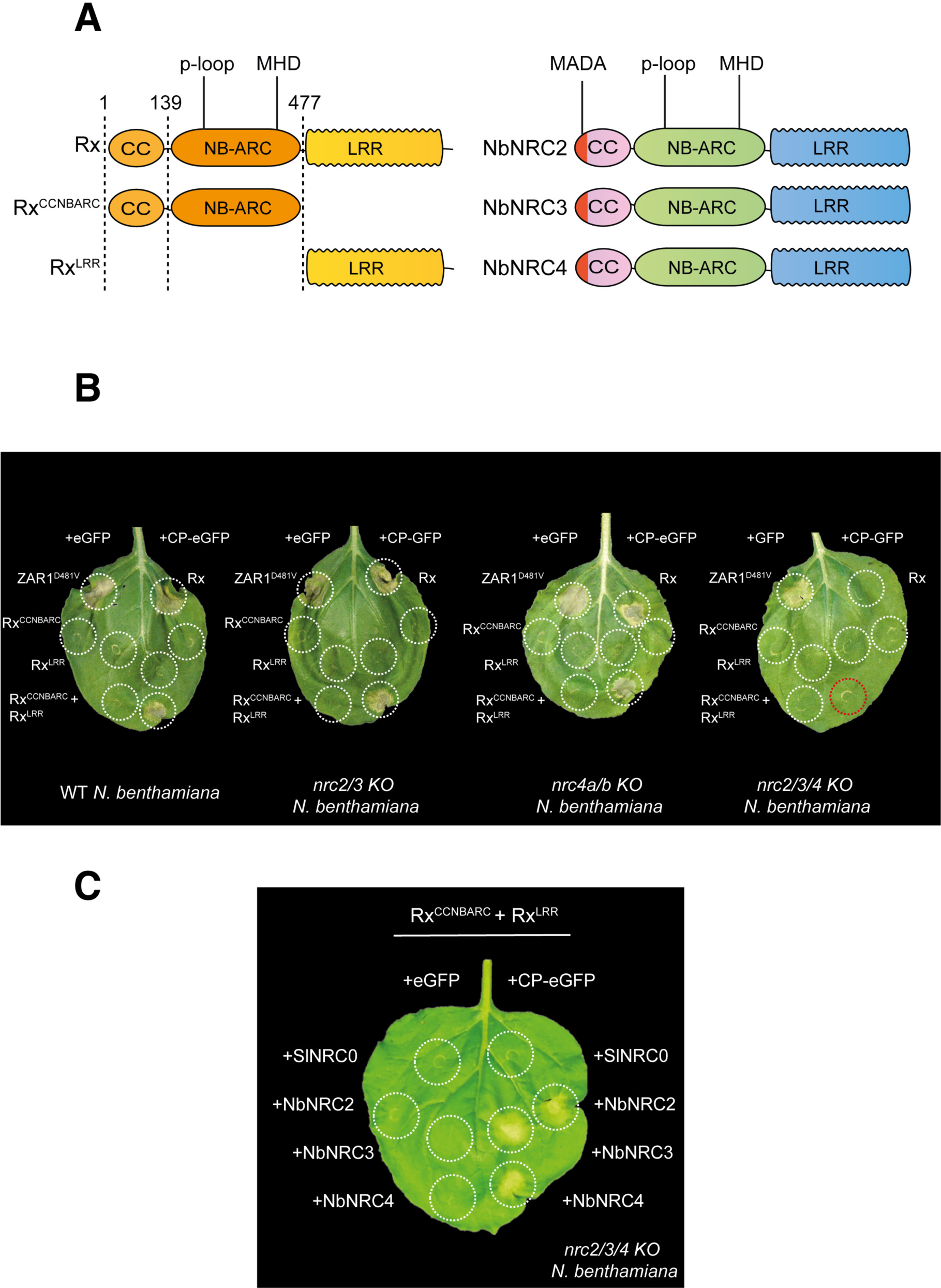
The cell death triggered by expression of Rx halves *in trans* upon perception of the coat protein (CP) of *Potato virus X* is dependent on NRC2, NRC3 or NRC4. (**A**) Schematic representation of Rx halves and NRC constructs used. Domain boundaries for Rx halves are indicated above. Approximate position of MADA, p-loop and MHD motifs are indicated above. Presence of an N-terminal MADA motif is indicated in dark red. (**B-C)** Representative leaves of different *N. benthamiana* KO lines agroinfiltrated to express constructs shown and photographed 5-7 days after infiltration. (**B**) Cell death mediated by Rx_CCNBARC_ and Rx_LRR_ complemented *in trans* is only abolished in *nrc2/3/4* KO *N. benthamiana* plants. Red dotted circle highlights absence of hypersensitive cell death in *nrc2/3/4* KO background. Wild-type Rx was included for comparison. NbZAR1_D481V_ was included as a control for NRC-independent cell death. (**C**) Cell death mediated by PVX CP-activated Rx_CCNBARC_ and Rx_LRR_ is complemented by NbNRC2, NbNRC3 and NbNRC4 in leaves of *nrc2/3/4* KO *N. benthamiana* lines when activated by co-expression of PVX CP. Free eGFP was used as a negative control for C-terminally eGFP-tagged PVX CP. SlNRC0 was used as a negative control as a helper NRC that does not get activated by Rx. Experiments were repeated 3 times with similar results.

### Rx halves mediate coat protein (CP)-dependent oligomerization of their NRC2 helper

We previously reported that Rx can mediate NRC2 oligomerization and resistosome formation upon PVX CP recognition, without forming part of the activated NRC2 oligomer (Contreras *et al*., 2023b; Contreras *et al*., 2023c). To test whether the activation of the Rx halves expressed *in trans* also leads to oligomerization of NRC2, we leveraged previously established blue-native polyacrylamide gel electrophoresis (BN-PAGE)-based readouts for NRC resistosome formation. We co-expressed the inactive or PVX CP-activated Rx^CCNBARC^ and Rx^LRR^ halves in leaves of *nrc2/3/4* KO *N. benthamiana* plants, complementing with the previously characterized NbNRC2^EEE^MADA motif mutant to abolish cell death induction without compromising helper activation (Contreras *et al*., 2023c). In these assays, Rx^CCNBARC^ and Rx^LRR^ expressed *in trans* mediated NRC2 oligomerization in a PVX CP-dependent manner, with both Rx halves being required. Notably, the NRC2 oligomer formed upon activation by Rx^CCNBARC^ and Rx^LRR^ migrates at the same height as the NRC2 oligomer formed upon activation by full-length Rx. The absence of a shift in size of the NRC2 oligomer in the Rx compared to the Rx^CCNBARC^/Rx^LRR^ treatments suggests that the PVX CP-activated Rx halves do not form part of the mature NRC2 oligomer, much like full-length Rx. (**Figure 2**).

**Figure 2:**
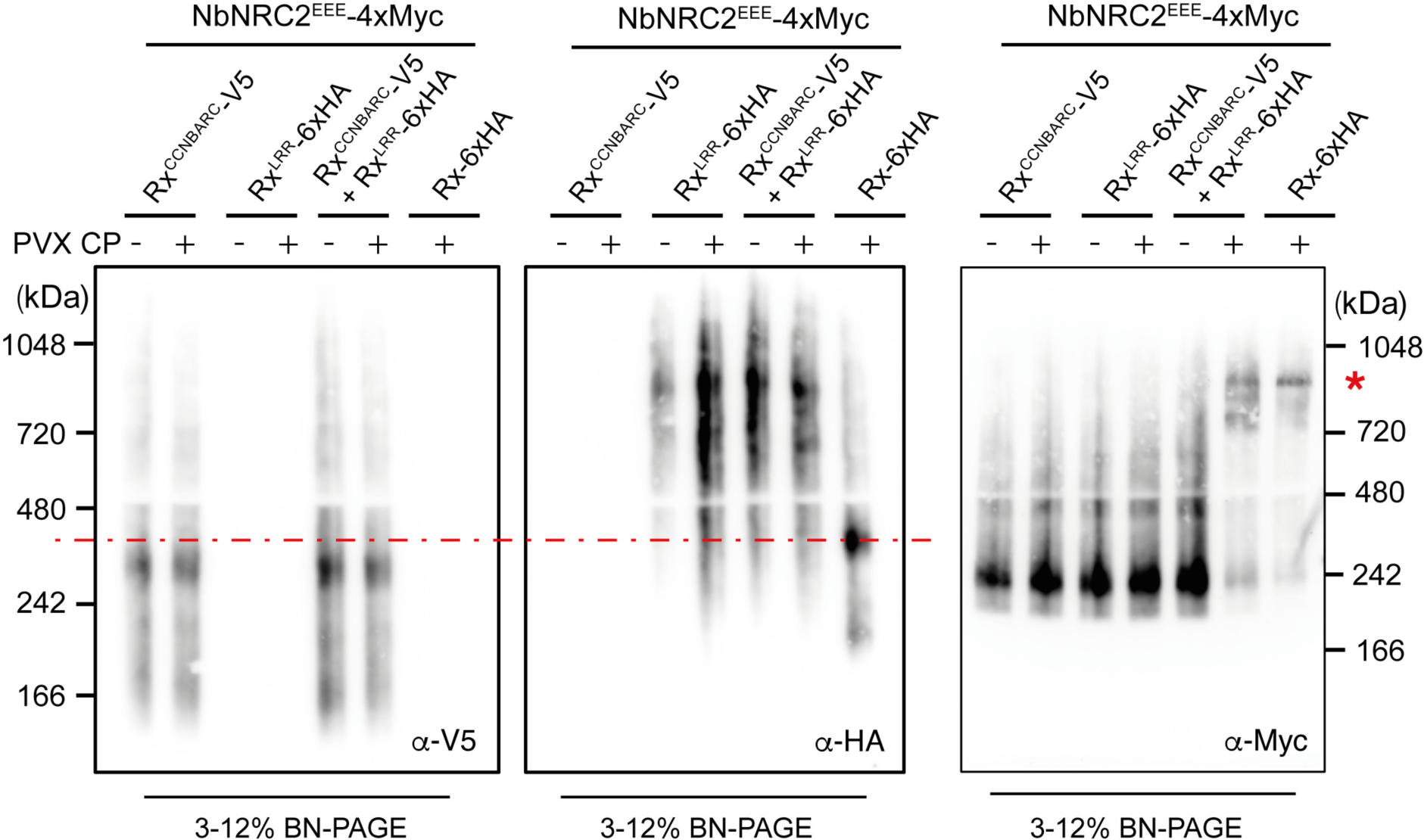
*Rx halves expressed in trans* mediate PVX CP-dependent oligomerization of their NRC2 helper. BN-PAGE assays with the inactive and PVX CP-activated Rx_CCNBARC_ and Rx_LRR_ halves co-expressed with NRC2_EEE_. C-terminally V5-tagged Rx_CCNBARC_, 6xHA-tagged Rx_LRR_ and 4xMyc-tagged NbNRC2_EEE_ were co-expressed with either free GFP or C-terminally GFP-tagged PVX CP. Full-length Rx was included for comparison and as a positive control for NRC2 oligomerization. Protein extracts were run on BN-PAGE and SDS-PAGE assays in parallel and immunoblotted with the appropriate antisera labelled on the bottom right corner of each blot. Red asterisks on the right indicates size of bands corresponding to the activated NRC2 complex. Red dotted lines indicate the molecular weight at which the full-length Rx complex migrates. V5 and HA blots were run on the same gel to allow for precise comparison of molecular weights. Approximate molecular weights (kDa) of the proteins are shown on the left for V5 and HA blots (run on the same gel), and on the right for Myc. SDS-PAGE accompanying BN-PAGE can be found in **Figure S1**. Experiments were repeated 3 times with similar results.

In previously published BN-PAGE assays, Rx is visualized as a complex of ∼ 400 kDa regardless of its activation state (Contreras & Kamoun, 2022; Contreras *et al*., 2023c). When probing for the Rx halves expressed *in trans* in BN-PAGE assays, we observed that Rx^CCNBARC^ also migrates as a band of ∼ 400 kDa, although this band migrates comparatively faster than full-length Rx indicating a lower molecular weight (**Figure 2**). Rx^LRR^, in contrast, is visualized as a high molecular weight smear covering molecular weights from around 400 to 1000 kDa. Like full-length Rx, we did not observe any change in the migration pattern of Rx^CCNBARC^ or Rx^LRR^ upon activation of the system with PVX CP. Although we can detect signal for Rx^LRR^ at a molecular weight that overlaps with the molecular weight of the NRC2 resistosome, this signal is also present in the treatments without PVX CP activation, suggesting that it is unlikely to correspond to a pool of Rx^LRR^ that is integrated into the NRC2 oligomer. We also noted that the migration pattern of Rx^CCNBARC^ is independent of the presence of Rx^LRR^, and vice-versa (**Figure 2**). We also did not observe a distinct band for Rx^LRR^ co-migrating with Rx^CCNBARC^ in BN-PAGE. The lack of co-migration in our BN-PAGE assays suggest that these two domains are not forming stable complexes and likely only associate transiently. Alternatively, a sub-pool of Rx halves that are forming a stable complex prior to activation are not highly abundant and cannot easily be detected in BN-PAGE.

### The effector-independent cell death triggered by the 154 amino acid nucleotide binding domain of Rx (Rx^NB^) is dependent on NRC2, NRC3 or NRC4

Previously, Rairdan and colleagues showed that a truncated version of the NRC-dependent sensor Rx encoding only the 154 amino acid NB domain region of the NB-ARC fused to eGFP (Rx^NB^-eGFP) was capable of constitutively triggering cell death in *N. benthamiana* and *N. tabacum* (Rairdan *et al*., 2008). Based on these results, we hypothesized that the NB domain could encode the minimal signal for NRC helper activation. To test this hypothesis, we performed cell death assays with Rx^NB^-eGFP in WT and *nrc2/3/4* KO *N. benthamiana* plants. We included the constitutively active ZAR1^D481V^ and Rx^D460V^ variants as controls for NRC-independent and NRC-dependent cell death, respectively, and eGFP as a negative control for cell death (**Figure 3A**). We were able to reproduce the previously reported cell death triggered by Rx^NB^-eGFP in WT *N. benthamiana*. This cell death was abolished in leaves of *nrc2/3/4* KO *N. benthamiana*, indicating that Rx^NB^-eGFP activates NRC-dependent hypersensitive cell death. This suggests that the NB domain of Rx is necessary and sufficient to activate downstream NRC helpers (**Figure 3A**).

**Figure 3:**
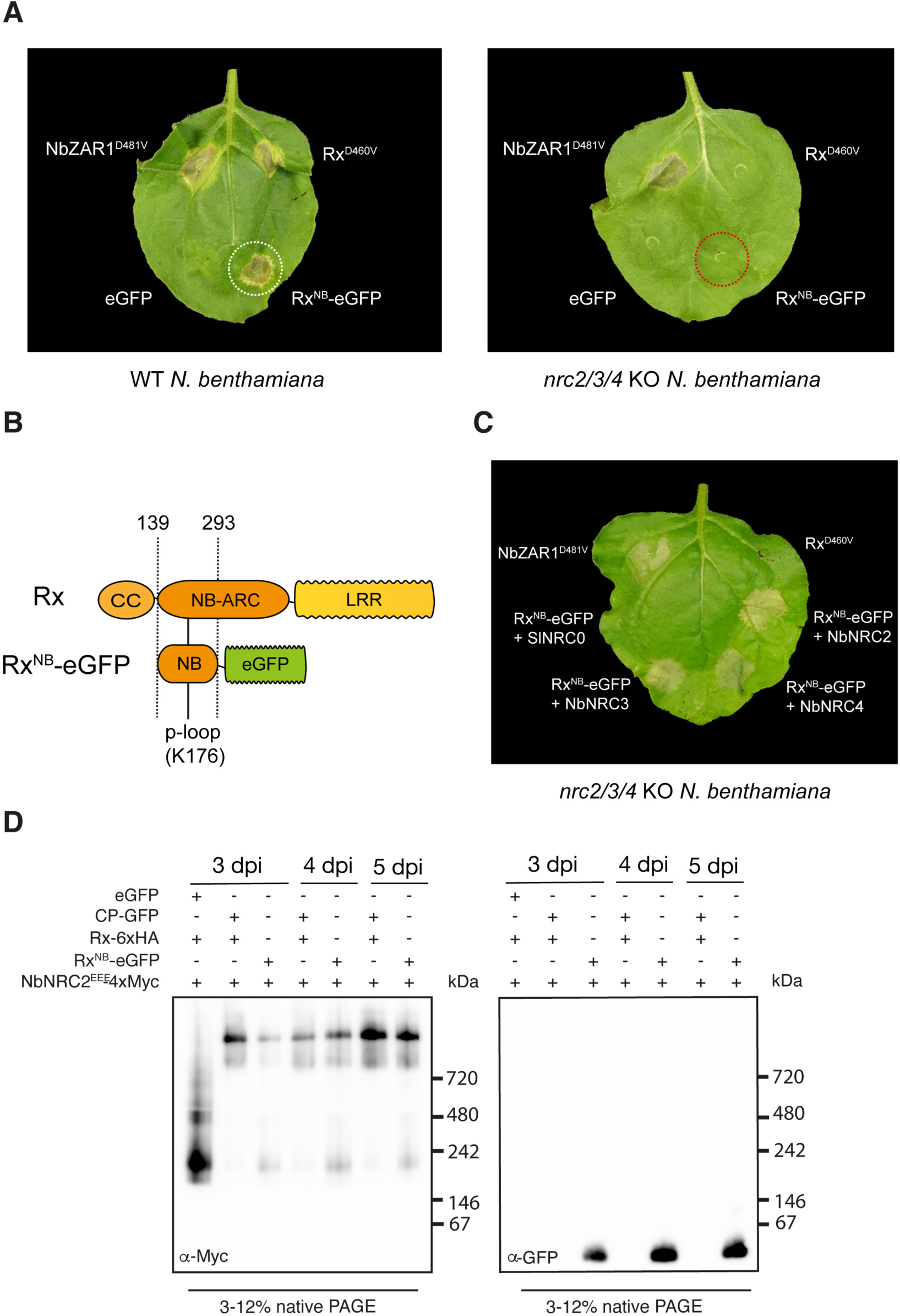
The NB domain of Rx (Rx^NB^) constitutively induces NRC2/NRC3/NRC4-dependent cell death and formation of the helper NRC2 oligomer. (**A**) Photo of a representative leaves from WT and *nrc2/3/4* KO *N. benthamiana* plants expressing Rx_NB_-eGFP. NbZAR1_D481V_ and Rx_D460V_ were included as controls for NRC-independent and NRC-dependent cell death, respectively. eGFP was included as a negative control for cell death. Leaves were agroinfiltrated to express the constructs indicated and photographed 5-7 days after infiltration. The experiment was repeated three times with at least 6 technical replicates per repeat, with similar results in all cases. (**B**) Schematic representation of Rx_NB_-eGFP construct. Domain boundaries used are indicated above, and were the same as reported previously by Rairdan and colleagues (Rairdan *et al*., 2008). Position of highly conserved lysine (K) residue of the p-loop is indicated, with numbering corresponding to its position in full-length Rx. (**C**) Representative leaf of *nrc2/3/4* KO *N. benthamiana* expressing Rx_NB_-eGFP complemented with NbNRC2, NbNRC3 or NbNRC4. SlNRC0 was included as a negative control for complementation. NbZAR1_D481V_ and Rx_D460V_ were included as controls for NRC-independent and NRC-dependent cell death, respectively. A quantitative analysis of the cell death can be found in **Figure S6**. (**D**) BN-PAGE assays with inactive and activated C-terminally 4xMyc-tagged NRC2_EEE_. NRC2_EEE_ was activated either with Rx-6xHA/PVX CP-eGFP or Rx_NB_-eGFP. Total protein extracts were run on native and denaturing PAGE assays in parallel and immunoblotted with the appropriate antisera labelled in the bottom left corner of each blot. Approximate molecular weights (kDa) of the proteins are shown on the right. Accompanying SDS-PAGE assays can be found in **Figure S4.** The experiment was repeated 3 times with similar results.

Mutating a highly conserved K residue in the p-loop motif typically makes NLRs non-functional (Mestre & Baulcombe, 2006). To test the p-loop dependency of Rx^NB^-eGFP-mediated activation of NRCs, we generated Rx^NB^-eGFP p-loop mutants (K to R substitution in position 176, as per numbering in full-length Rx, **Figure 3B**) and expressed them in leaves of WT and *nrc2/3/4* KO *N. benthamiana*. Interestingly, much like Rx^NB^-eGFP, the Rx^NB^-eGFP p-loop mutants triggered NRC-dependent cell death. This indicates that unlike full-length Rx and the Rx halves expressed *in trans* (Moffett *et al*., 2002), cell death triggered by Rx^NB^-eGFP does not require an intact p-loop (**Figure S2**).

To further test the NRC-dependency of Rx^NB^-eGFP triggered cell death, we carried out complementation assays in leaves of *nrc2/3/4* KO *N. benthamiana*. We co-expressed Rx^NB^-eGFP with NbNRC2, NbNRC3 and NbNRC4 or SlNRC0, a tomato NRC that full-length Rx is unable to signal through (Sakai *et al*., 2023). The cell death mediated by the Rx^NB^-eGFP was restored upon complementation with NbNRC2, NbNRC3 and NbNRC4, but not SlNRC0 (**Figure 3C**). We conclude that Rx^NB^-eGFP is indeed activating cell death via NRC-dependent pathways and that it retains the specificity to activate the same set of NRC helpers that full-length Rx signals through in *N. benthamiana*.

To investigate the contributions of the C-terminal eGFP tag to the cell death mediated by Rx^NB^-eGFP, we generated Rx^NB^-mCherry-6xHA fusions and tested them for NRC-dependent cell death. These Rx^NB^-mCherry-6xHA fusions also triggered NRC-dependent cell death, although this cell death was weaker than that triggered by Rx^NB^-eGFP (**Figure S3A**). In complementation assays carried out in leaves of *nrc2/3/4* KO *N. benthamiana*, cell death triggered by Rx^NB^-mCherry-6xHA was also complemented by NbNRC2, NbNRC3 and NbNRC4 but not SlNRC0 (**Figure S3B**). We used western blot analysis to test for Rx^NB^-mCherry-6xHA protein accumulation levels in *nrc2/3/4* KO *N. benthamiana*. Strikingly, although this construct triggers NRC-dependent cell death in WT *N. benthamiana*, we were unable to detect any signal for Rx^NB^-mCherry-6xHA (**Figure S3C**). Based on these results, we conclude that the cell death triggered by the Rx^NB^ truncations is not specific to Rx^NB^-eGFP. Since Rx^NB^-eGFP-triggered cell death is much stronger and easier to detect in planta, we decided to proceed with C-terminally eGFP-tagged variants for all subsequent experiments.

### Rx^NB^ mediates effector-independent oligomerization of the NRC2 helper

We next sought to investigate if Rx^NB^ is sufficient to trigger NRC2 oligomerization. To this end, we co-expressed Rx^NB^ and NbNRC2^EEE^ in leaves of *nrc2/3/4* KO *N. benthamiana* and performed BN-PAGE assays at different timepoints, with Rx/PVX CP as a positive control for NRC2 oligomerization. Interestingly, expression of Rx^NB^-eGFP was sufficient to mediate the formation of NRC2 high molecular weight complexes of a similar size as the NRC2 oligomers formed upon activation with full-length Rx/PVX CP. Importantly, the absence of a shift in size of the NRC2 oligomers triggered by Rx^NB^-eGFP compared to those triggered by Rx/PVX CP suggests that, much like full-length Rx, Rx^NB^-eGFP does not form part of the mature NRC2 oligomer (**Figure 3D**). We noted that compared to the NRC2 oligomer signal induced by full-length Rx and PVX CP, Rx^NB^-eGFP-activated NRC2 treatments showed a weak signal corresponding to inactive NRC2 in all timepoints analyzed, indicating that Rx^NB^-eGFP mediated helper oligomerization may be less efficient (**Figure 3D**). We did not observe any signal for Rx^NB^-eGFP at a molecular weight matching that of the NRC2 oligomer, which further suggests that the mature NRC2 resistosome does not include Rx^NB^-eGFP (**Figure 3D**).

### The nucleotide binding (NB) domains of the sensor NLRs Gpa2, Rpi-amr1e, Rpi-amr3 and Sw-5b are sufficient to activate downstream NRC helpers

To what degree does the activation of NRC helpers by the NB domain of sensor NLRs apply to other disease resistance proteins? We investigated this by generating NB domain-eGFP fusions of 8 well-studied disease resistance NLR proteins that are dependent on NRC helpers in the immune receptor network (**Figure 4A, Figure S5**). This panel covered the phylogenetic diversity of NRC-dependent sensors and was composed of the Rx-type sensors Gpa2, Rpi-amr1e, Rpi-amr3 and Bs2, and the SD-type sensors Mi-1.2, Rpi-blb2 and Sw-5b (**Figure 4A**) (Contreras *et al*., 2023a). Gpa2^NB^-eGFP, Rpi-amr1e^NB^-eGFP and Sw-5b^NB^-eGFP triggered NRC-dependent cell death to levels comparable to Rx^NB^-eGFP (**Figure 4B, Figure S6**). Rpi-amr3^NB^-eGFP triggered weak NRC-dependent cell death in a subset of the leaves tested (6 out of 18 total infiltration spots) (**Figure 4B**, **Figure S6**). In contrast, Bs2^NB^-eGFP, Rpi-blb2^NB^-eGFP and Mi-1.2^NB^-eGFP did not trigger cell death (**Figure 4B, Figure S6)**. In parallel, we determined the protein accumulation levels of each NB domain-eGFP fusion *in planta*. All of the sensor NLR NB domains that induced cell death accumulated *in planta* (Rx, Gpa2, Rpi-amr1e and Sw-5b). In contrast, the NB domain truncations that triggered weak or no cell death accumulated to comparatively lower levels (Rpi-amr3, Bs2, Rpi-blb2 and Mi-1.2) (**Figure S7**). We conclude that NB domain-mediated activation of downstream NRC helpers is not exclusive to Rx and can also be triggered by other Rx-type and SD-type sensors of the NRC network.

**Figure 4:**
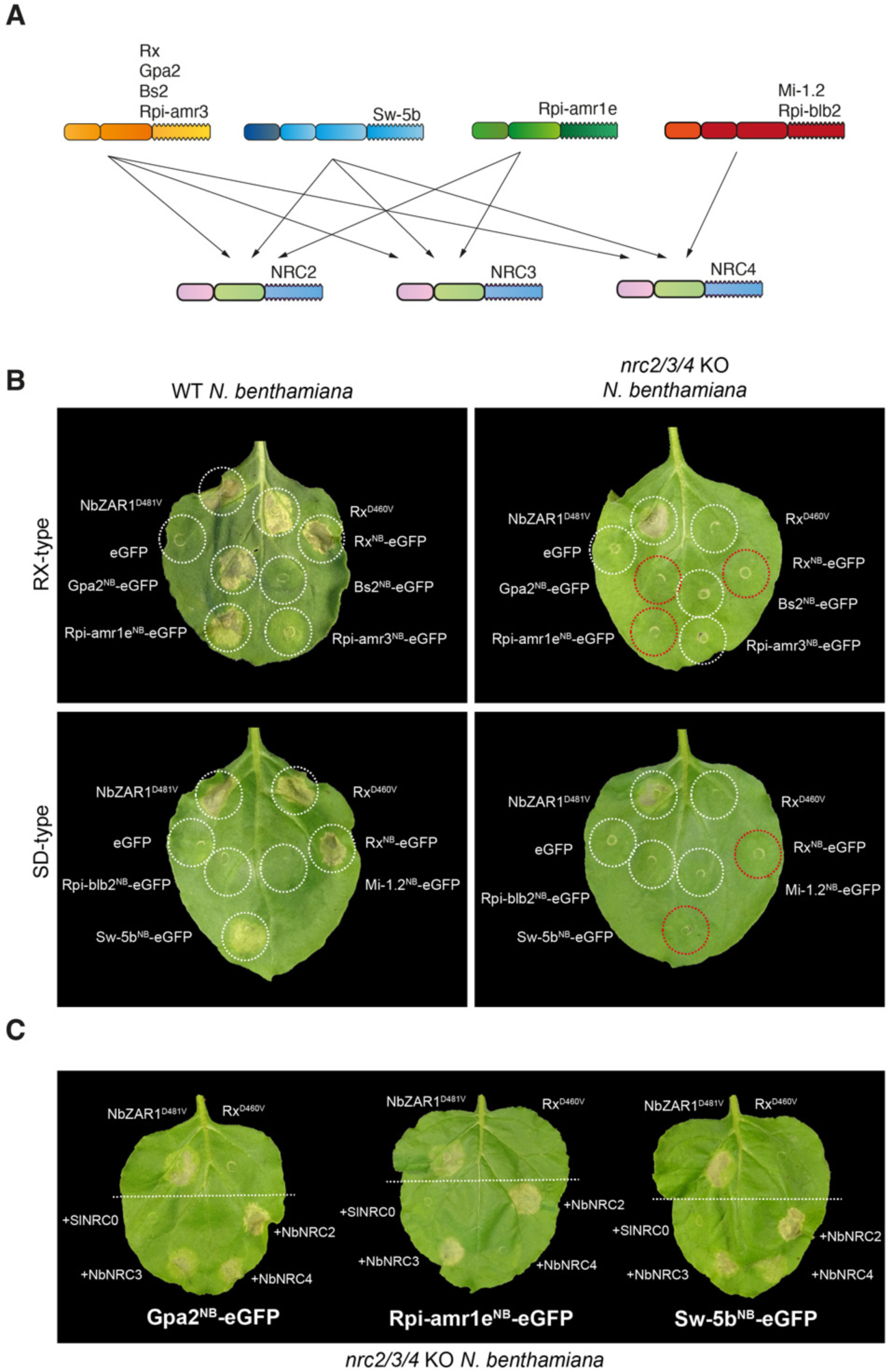
The NB domains of the sensor NLRs Gpa2, Rpi-amr1e, Rpi-amr3 and Sw-5b are sufficient to activate downstream NRC helpers. (**A**) Schematic representation of sensor-helper signaling specificities in the NRC network. Rx-type sensors Rx, Gpa2, Rpi-amr1e and Rpi-amr3 are represented with three domains (CC, NB-ARC and LRR) whereas SD-type sensors Sw-5b, Rpi-blb2 and Mi-1.2 are represented with four domains (SD, CC, NB-ARC and LRR). (**B**) Representative photos of cell death assays with the constructs indicated in leaves of either WT or *nrc2/3/4* KO *N. benthamiana*. NbZAR1_D481V_ and eGFP were included as positive and negative controls for cell death, respectively. Rx_D460V_ was included as a control for NRC-dependent cell death. Leaves were agroinfiltrated to express the constructs indicated and photographed 5-7 days after infiltration. One representative leaf is shown. A quantitative analysis of the cell death assays can be found in **Figure S6.** (**C**) Representative photos of cell death assays with the constructs indicated in leaves of *nrc2/3/4* KO *N. benthamiana*. NbZAR1_D481V_ and Rx_D460V_ were included as positive and negative controls for cell death, respectively. Leaves were agroinfiltrated to express the constructs indicated and photographed 5-7 days after infiltration. One representative leaf is shown. A quantitative analysis of the cell death assays can be found in **Figure S6**.

### The nucleotide binding (NB) domains of the sensor NLRs Gpa2, Rpi-amr1e and Sw-5b retain the NRC helper specificities of their full-length counterparts

Sensor NLRs in the NRC network can signal via different subsets of downstream NRC helpers. For example, PVX CP-activated Rx, RBP1-activated Gpa2 and NSm-activated Sw-5b can signal interchangeably via NRC2, NRC3 and NRC4, whereas AVRamr1-activated Rpi-amr1e can signal via NRC2 and NRC3 but not NRC4 (**Figure 4A, Figure S8**) (Contreras *et al*., 2023b; Contreras *et al*., 2023c; Derevnina *et al*., 2021; Lin *et al*., 2022; Witek *et al*., 2021; Wu *et al*., 2017). We next sought to determine whether, Gpa2^NB^-eGFP, Rpi-amr1e^NB^-eGFP and Sw-5b^NB^-eGFP retained the same downstream NRC specificities of their full-length counterparts, similar to Rx^NB^-eGFP. To this end, we performed complementation assays in leaves of *nrc2/3/4* KO *N. benthamiana* co-expressing each of these NB domains together with NbNRC2, NbNRC3 or NbNRC4, respectively. We included SlNRC0 as a negative control for complementation. We did not include Rpi-amr3 in these experiments as Rpi-amr3^NB^-eGFP accumulated poorly *in planta* and triggered comparatively weaker cell death, making it harder to draw robust conclusions (**Figure 4**, **Figure S6, Figure S7**). The cell death triggered by Gpa2^NB^-eGFP and Sw-5b^NB^-eGFP was restored upon co-expression with NbNRC2, NbNRC3 and NbNRC4. Rpi-amr1e^NB^-eGFP cell death was restored upon co-expression with NbNRC2 and NbNRC3 but not with NbNRC4. (**Figure 4C**). None of the NB domains was complemented by SlNRC0. This indicates that NB domain-eGFP fusions retain the same helper NRC specificity profiles of their full-length counterparts.

### Rx^NB^ and its helper NRC2 form a minimal functional unit that can be transferred from Solanaceae to the Asteraceae plant lettuce

We leveraged transient expression in lettuce (*Lactuca sativa*) by agroinfiltration (Wróblewski *et al*, 2018) to investigate the capacity of the Rx/NRC2 pair to function in a species distantly related to the Solanaceae. Lettuce is an Asteraceae in the Campanulid lineage and is estimated to have split from *Nicotiana* (Solanaceae, Lamiid) about 102 Million years ago (Zeng *et al*, 2014). Lettuce predates the expansion of the NRC helpers that occurred in Solanaceae and other lamiids and therefore lacks orthologs of NRC2, NRC3 and NRC4 and other Solanaceae NRCs (Goh *et al*., 2023; Sakai *et al*., 2023). We first determined whether Rx/CP and NRC2 can trigger cell death in lettuce by co-expressing these three proteins in lettuce leaves. We included an autoactive variant of the previously published lettuce CC_G10_-NLR RGC2B (RGC2B^D470V^) as a positive control for cell death (Meyers *et al*, 1998; Shen *et al*, 2002). We observed a cell death response indicative of immune activation when all three components, CP, Rx and NRC2 were co-expressed in leaves of lettuce. This cell death was abolished when any one of these three components was missing (**Figure 5A**).

**Figure 5:**
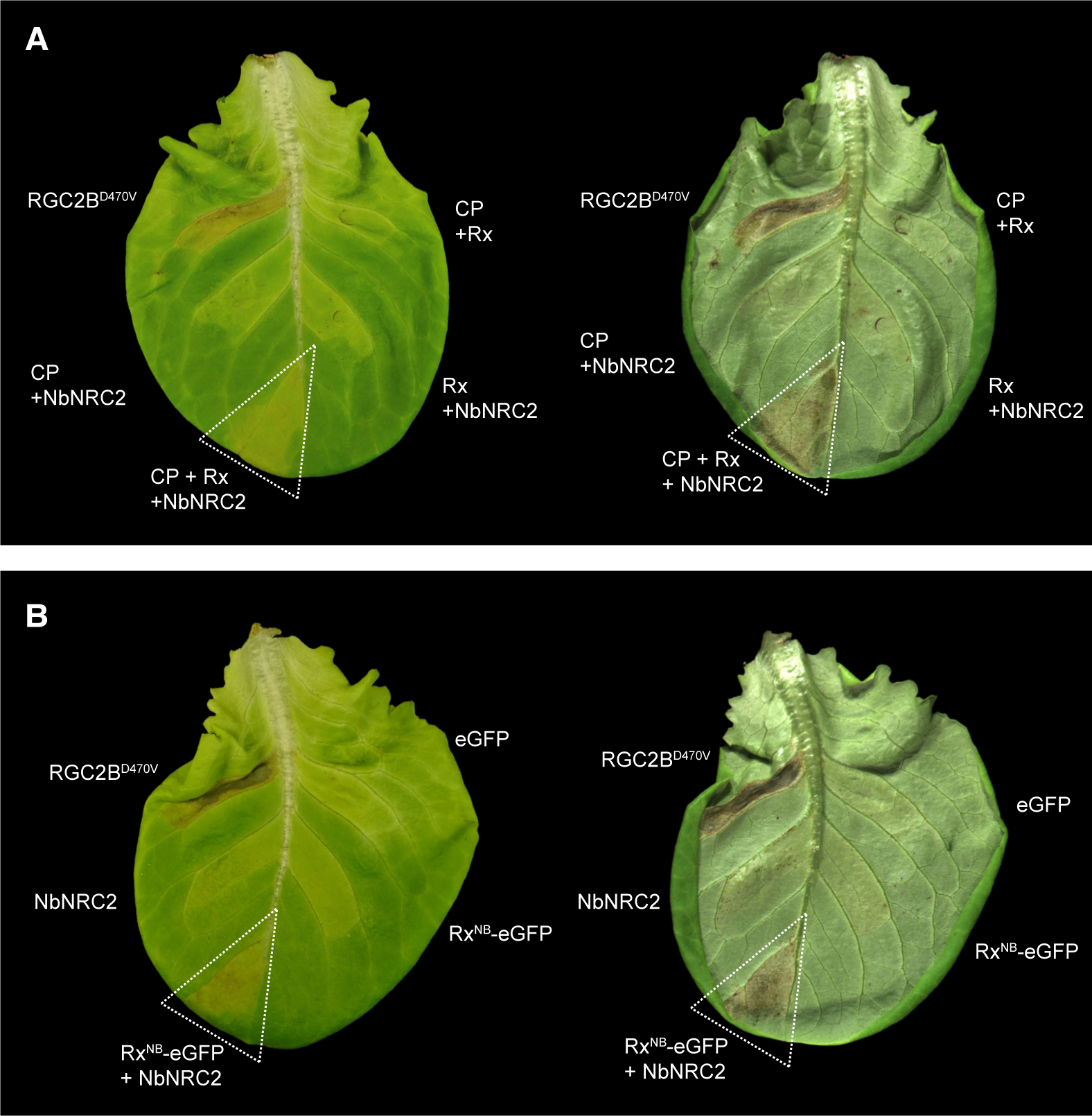
Rx, Rx^NB^ and their helper NRC2 form a minimal functional unit that can be transferred to the Asteraceae plant lettuce. Photo of representative leaves of lettuce (*L. sativa*) var. “Fenston”. (**A**) Lettuce leaf showing cell death after co-expression of PVX CP, Rx and NbNRC2. No cell death was observed when any one of the three components (PVX CP, Rx or NbNRC2) was absent. (**B**) Photo of representative leaves of lettuce (*L. sativa*) var. “Fenston” showing cell death after co-expression of Rx_NB_-eGFP and NbNRC2. No cell death was observed with eGFP or with Rx_NB_-eGFP and NbNRC2 when expressed individually. (**A-B**) Images were taken 5-7 days after agroinfiltration, and leaves were imaged from both adaxial (left side of panel) and abaxial (right side of panel) sides. Images from abaxial side were included as HR cell death in lettuce is more visible on abaxial than adaxial sides. The constitutively active variant of the lettuce NLR RGC2B (RGC2B_D470V_) was used as a positive control for cell death in lettuce. Experiments were repeated three times with similar results.

Next, we tested whether Rx^NB^-eGFP and NRC2 were also capable of mediating cell death in lettuce. We included RGC2B^D470V^ and eGFP as positive and negative controls for cell death, respectively. Co-expression of Rx^NB^-eGFP and NbNRC2 triggered cell death in lettuce, whereas expression of either Rx^NB^-eGFP or NbNRC2 alone did not (**Figure 5B**). This result suggests that no additional proteins are required for Rx^NB^-eGFP to activate NRC2. Alternatively, additional proteins that may be required are conserved between *N. benthamiana* (Lamiid) and lettuce (Campanulid). We conclude that Rx^NB^-eGFP and NRC2 are most likely a minimal two-component system that can be transferred from *N. benthamiana* to lettuce.

## Discussion

In this study, we build upon findings by Moffett, Rairdan, and colleagues regarding the ability of individual domains of the virus disease resistance protein Rx to activate immunity *in trans*, or as a nucleotide-binding (NB) domain (Moffett *et al*., 2002; Rairdan *et al*., 2008). We revisited this work within the framework of the activation-and-release biochemical model for sensor-helper activation in the NRC network of CC-NLRs (Contreras *et al*., 2023c). Our findings demonstrates that PVX CP-triggered cell death, mediated by the CC-NBARC (Rx^CCNBARC^) and LRR (Rx^LRR^) domains of Rx when expressed *in trans*, as well as the effector-independent cell death mediated by Rx^NB^-eGFP, are dependent on NRC helpers and involves the formation of NRC2 resistosomes. Similar to the full-length Rx, the Rx halves and Rx^NB^-eGFP do not integrate into the mature NRC2 helper oligomer (**Figure 1**, **Figure 2**, **Figure 3**). Furthermore, our work reveals that the ∼150 amino acid NB domain fragments of other sensor NLRs, such as Gpa2, Rpi-amr1e, Rpi-amr3 and Sw-5b, also induce constitutive cell death. Importantly, Rx^NB^, Gpa2^NB^, Rpi-amr1e^NB^, and Sw-5b^NB^ maintain the downstream helper specificity profiles of their full-length counterparts (**Figure 3**, **Figure 4**). Finally, we demonstrate that both full-length PVX CP-activated Rx and Rx^NB^-eGFP can initiate cell death in lettuce when co-expressed with NRC2, suggesting that this represents a minimal, two-component system transferable to unrelated plant taxa (**Figure 5**). Our data support a working model in which the conditional exposure of sensor NLR NB domains upon pathogen perception serves as a signal for downstream NRCs that triggers helper oligomerization and resistosome formation (**Figure 6**).

**Figure 6:**
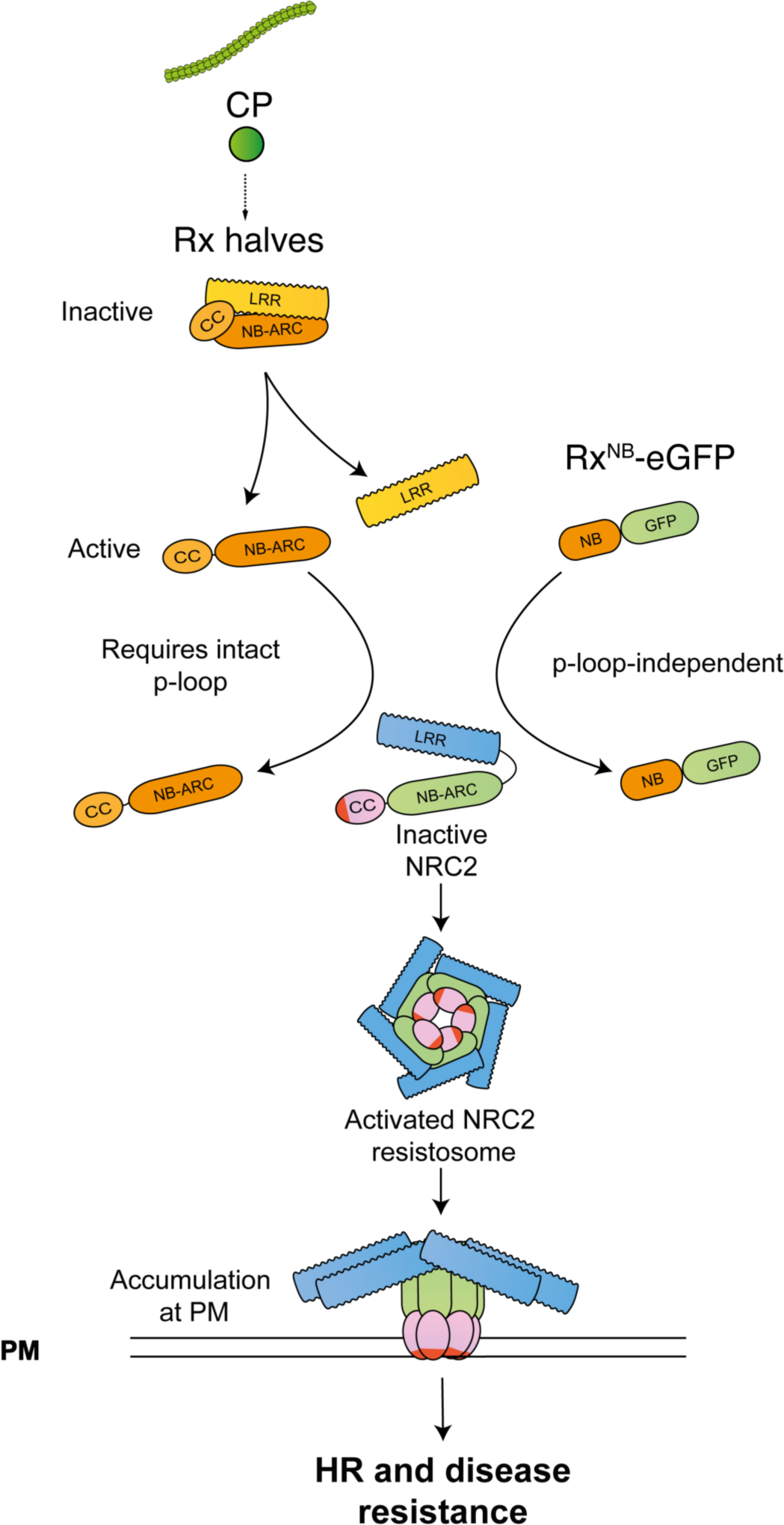
Proposed working model for NRC activation by Rx halves expressed *in trans* and Rx^NB^-eGFP. Schematic representation of Rx halves and Rx_NB_-eGFP-mediated NRC activation. Prior to effector-triggered activation, NRC-dependent sensors such as Rx are held in an inactive conformation by intramolecular interactions. These intramolecular interactions between the different Rx domains hide the sensor NLR NB domain, preventing its signaling, and contribute to the previously reported association of the Rx_CCNBARC_ and Rx_LRR_ halves (Moffett *et al*., 2002). Upon perception of PVX CP, Rx undergoes a series of p-loop dependent conformational changes that expose the NB domain and lead to dissociation of the Rx halves expressed *in trans* (Moffett *et al*., 2002). This conditional NB domain exposure allows it to signal to downstream helpers such as NRC2 leading to its homo-oligomerization and resistosome formation. Following activation by their upstream sensor, NRC helper oligomers signal via their N-terminal CC domain. In the case of the Rx_NB_-eGFP and other sensor NB domain-eGFP fusions, the NB domain is exposed and therefore constitutively activates downstream helpers. Because no intramolecular rearrangements are required to relieve autoinhibitory intramolecular interactions, the p-loop mutation does not abolish this cell death.

The prevailing paradigm for NLR activation is that their N-terminal domains mediate immune signaling upon resistosome assembly. This is the case for the N-terminal CC domains of singleton and helper CC-NLRs and for the N-terminal TIR domains of sensor TIR-NLRs (Bentham *et al*., 2018; Contreras *et al*., 2023a). Based on our data, we propose that for NRC dependent sensor NLRs, the NB domain region of the central NB-ARC, not the N-terminal CC domain, executes signaling to downstream helpers. How does the NB domain of sensor NLRs initiate the activation of NRC helpers? Is this activation achieved through direct binding or does it necessitate enzymatic activity, as is the case for TIR-NLRs and their downstream CC_R_-NLR helpers (Huang *et al*, 2022; Ma *et al*, 2020)? The activation of full-length Rx and the Rx halves expressed *in trans* requires an intact p-loop (Moffett et al., 2002). In sharp contrast, Rx^NB^ activation of NRCs is p-loop independent (**Figure S2**). This observation indicates that while ATP-binding is required for conditional NB domain exposure, it does not play a role in the recognition of the sensor NB domain by the helper once the NB domain is exposed, making sensor NB domain-catalyzed ATP hydrolysis an improbable mechanism of helper activation.

Our transient expression assays in lettuce, along with recently published data demonstrating that Campanulid NRC sensor-helper pairs can initiate immune signaling in *N. benthamiana* (Goh et al., 2023; Sakai et al., 2023), lead us to favor a model in which sensor NB domains and NRC helpers directly interact, with no additional components required for sensor-helper communication. While it remains possible that additional conserved components exist between *N. benthamiana* and lettuce, the evolutionary divergence between these two plant species—estimated at 102 million years— makes this hypothesis less likely (Zeng *et al*., 2014). Importantly, BN-PAGE assays do not reveal any evidence of NB domain integration into the NRC2 resistosome, suggesting that this interaction is transient, as previously suggested for full-length sensor activation of NRCs (Contreras *et al*., 2023c). To date, conclusive evidence that NRC-dependent sensors and their NRC helpers form stable complexes has not been obtained, possibly because of the transient nature of this interaction. Notably, that the NB domain alone is sufficient to activate NRCs suggests that sensor NLRs are unlikely to be required as scaffolds for helper NLR polymerization during sensor-helper activation, as is the case for the mammalian NAIP/NLRC4 sensor-helper pairs (Zhang *et al*, 2015). Further biochemical and structural analyses will provide insights into the precise dynamics of sensor-helper interactions during activation.

According to a well-established paradigm of plant immunity, NLRs are known to guard critical host components, known as guardees or decoys, that are perturbed during pathogen infection (Dangl & Jones, 2001; Van Der Biezen & Jones, 1998; van der Hoorn & Kamoun, 2008). We propose that helper NLRs, such as the NRCs, may have evolved to guard the conformational state of the NB-ARC domains in their upstream sensors, enabling them to detect any pathogen-triggered modifications. The NB-ARC domain stands out as the most conserved feature within the NLR protein family, and NB-ARC rearrangements have been demonstrated as critical for NLR activation across various life forms (Chou *et al*., 2023; Duxbury *et al*., 2021). It is plausible that NRC helpers have evolved to monitor the activation status of the NB-ARC domains of their upstream sensors, recognizing NB-ARC conformational states indicative of effector perception and immune activation. This concept is reminiscent of how CC_R_-NLR helpers, such as NRG1 and ADR1, have evolved to sense the activation of their upstream TIR-NLR sensors by detecting the formation of EDS1-SAG101/PAD4 heterodimers induced following TIR-NLR resistosome assembly and small molecule production (Feehan et al., 2023; Huang et al., 2022; Jia et al., 2022). We propose that helper NLRs in the NRC family function as “guards of sensors”, monitoring sensor NLR activation. In this model, sensor NLRs could even be considered as decoys, as they have degenerated features such as non-functional MADA motifs and N-terminal domain integrations (SD) and have lost the capacity to trigger cell death on their own (Adachi *et al*., 2019b; Contreras *et al*., 2023a).

The discovery that Rx and NRC2 can effectively function in lettuce carries significant implications for disease resistance breeding and bioengineering (Marchal *et al*, 2022). These findings raise the possibility of transferring NRC sensor-helper pairs across crop species at least within Asterid plants. This potential transfer may even extend to agronomically important species outside of the Asteraceae, such as soybean, wheat, or rice, which lack NRCs altogether (Wu *et al*., 2017). Furthermore, it opens the door to the prospect of transferring entire networks of NRC sensors and helpers across species boundaries, capitalizing on the resilience that networked signaling architectures offer to immune systems (Wu *et al*., 2018). The transfer of the NRC network across species is particularly exciting given that it confers resistance to multiple species of pathogens and pests (Derevnina *et al*., 2021; Kourelis *et al*., 2022; Wu *et al*., 2017). Additionally, the observation that Rx can recognize PVX CP in lettuce, triggering cell death, prompts important inquiries about the recognition mechanism of PVX CP. Previous work suggested that Rx indirectly recognizes PVX CP and that this recognition necessitates the host component RanGAP2 (Moffett et al., 2002; Sacco et al., 2007; Tameling & Baulcombe, 2007). The fact that Rx can still respond to PVX CP in lettuce raises the possibility that either Rx retains its proper functionality with the lettuce RanGAP2 ortholog or that Rx recognition of PVX CP might involve direct, albeit transient, interactions. Further research is required to elucidate the precise role of RanGAP2 in Rx functionality and to unravel the specific mechanism by which Rx perceives PVX CP.

A significant, unresolved question is deciphering the molecular basis of sensor-helper specificity, a fundamental characteristic of the NRC network. While certain sensors can activate multiple helpers within a given host, others selectively activate a subset of the NRC helper repertoire. Furthermore, across evolutionary timescales, sensors from one species can lose the ability to communicate with NRC helpers from distantly or even closely related species, as is the case of species-specific sensor-helper NRC pairs in lamiids (Goh *et al*., 2023; Sakai *et al*., 2023). Our data suggests that the NB domain plays a crucial role as a molecular determinant for sensor-helper specificity on the sensor side. However, although the sensor NB is necessary and sufficient for NRC activation, the contributions of other sensor NLR domains to downstream helper activation remain unclear. Furthermore, the domains within the helper proteins involved in monitoring sensor NLR activation are yet to be identified. Understanding the domains and interfaces involved in sensor and helper protein interactions, and their evolutionary dynamics, will provide additional insights into the mechanisms through which paired and networked CC-NLRs activate. Identifying the precise molecular communication between sensors and helpers could have substantial implications for disease resistance engineering, potentially enabling the synthetic expansion and optimization of sensor-helper interactions, resulting in more robust and efficient immune receptor networks.

## Materials and Methods

### Plant growth conditions

Wild-type, *nrc2/3*, *nrc4a/b* and *nrc2/3/4* KO *Nicotiana benthamiana* lines as well as lettuce (*L. sativa*) var. “Fenston” were grown in a controlled environment growth chamber with a temperature range of 22–25°C, humidity of 45–65% and a 16/8-h light/dark cycle.

### Plasmid construction

The Golden Gate Modular Cloning (MoClo) kit (Weber *et al*, 2011) and the MoClo plant parts kit (Engler *et al*, 2014) were used for cloning, and all vectors are from this kit unless specified otherwise. Cloning design and sequence analysis were done using Geneious Prime (v2021.2.2; https://www.geneious.com). Rx, Rx^D460V^, CP-eGFP and NRC2^EEE^ constructs used were previously described (Contreras *et al*., 2023c). NbZAR1^D481V^ construct was previously described (Harant *et al*., 2022). All NRC constructs used were previously described (Derevnina *et al*., 2021; Sakai *et al*., 2023). Rx halves were cloned into pICH86988 acceptor with integrated 35S promoter and OCS terminator. All NB domain constructs were cloned into the pJK268c vector with pICSL51288 (2×35S promoter) and pICSL41414 (35S terminator). Domain boundaries for all sensor NB domains were based on amino acid sequence alignment to the original Rx^NB^ domain boundaries reported by Rairdan and colleagues (**Figure S5**) (Rairdan *et al*., 2008), and subsequently ordered as synthetic gene fragments (Azenta/Genewiz). RGC2B^D470V^ construct was synthesized and cloned into pJK001c acceptor with pICSL51288 (2×35S promoter) and pICSL41414 (35S terminator). C-terminal tag modules used were pICSL50012 (V5), pICSL5007 (3xFLAG), pICSL50034 (eGFP), pICSL5009 (6xHA) or pICSL50010 (4xMyc), as indicated.

### Cell death assays by agroinfiltration

Proteins of interest were transiently expressed in *N. benthamiana* and lettuce according to previously described methods (Bos et al, 2006). Briefly, leaves from 4–5-week-old plants were infiltrated with suspensions of *A. tumefaciens*, using the GV3101 pM90 strain for *N. benthamiana* and the C58C1 strain for lettuce. Strains were transformed with expression vectors coding for different proteins indicated. Final OD_600_ of all *A. tumefaciens* suspensions were adjusted in infiltration buffer (10 mM MES, 10 mM MgCl2, and 150 μM acetosyringone (pH 5.6)). Final OD_600_ used was 0.3 for each construct. Whenever multiple constructs were co-infiltrated into an individual spot, the total concentration of bacteria was kept constant across treatments by adding untransformed *A. tumefaciens* when necessary. This was to avoid an effect from differences in the total OD_600_ of bacteria in each treatment.

### Extraction of total proteins for BN-PAGE and SDS–PAGE assays

Four to five-week-old plants were agroinfiltrated as described above with constructs of interest and leaf tissue was collected 3 days post agroinfiltration. Final OD_600_ used was 0.3 for each NLR immune receptor, Rx halves or NB domain, and 0.2 for eGFP or CP-eGFP. BN-PAGE was performed using the Bis-Tris Native PAGE system (Invitrogen) according to the manufacturer’s instructions. Leaf tissue was ground using a Geno/Grinder tissue homogenizer and total protein was subsequently extracted and homogenized extraction buffer. GTMN extraction buffer was used (10% glycerol, 50 mM Tris–HCl (pH 7.5), 5 mM MgCl_2_ and 50 mM NaCl), supplemented with 10 mM DTT, 1x protease inhibitor cocktail (SIGMA) and 0.2% Triton X-100 (SIGMA). Samples were incubated in extraction buffer on ice for 10 min with short vortex mixing every 2 min. Following incubation, samples were centrifuged at 5,000x*g* for 15 min and the supernatant was used for BN-PAGE and SDS–PAGE assays.

### BN-PAGE assays

BN-PAGE assays were done as described previously (Ahn *et al*., 2023; Contreras *et al*., 2023c). Brieflly samples extracted as detailed above were diluted as per the manufacturer’s instructions by adding NativePAGE 5% G-250 sample additive, 4x Sample Buffer and water. After dilution, samples were loaded and run on Native PAGE 3–12% Bis-Tris gels alongside either NativeMark unstained protein standard (Invitrogen) or SERVA Native Marker (SERVA). The proteins were then transferred to polyvinylidene difluoride membranes using NuPAGE Transfer Buffer using a Trans-Blot Turbo Transfer System (Bio-Rad) as per the manufacturer’s instructions. Proteins were fixed to the membranes by incubating with 8% acetic acid for 15 min, washed with water and left to dry. Membranes were subsequently re-activated with methanol to correctly visualize the unstained native protein marker. Membranes were immunoblotted as described below.

### SDS–PAGE assays

For SDS–PAGE, samples were diluted using a 3:1 ratio of sample to SDS loading dye and denatured at 72 °C for 10 min. Denatured samples were spun down at 5,000 *g* for 3 min and supernatant was run on 4–20% Bio-Rad 4–20% Mini-PROTEAN TGX gels alongside a PageRuler Plus prestained protein ladder (Thermo Scientific). The proteins were then transferred to polyvinylidene difluoride membranes using Trans-Blot Turbo Transfer Buffer using a Trans-Blot Turbo Transfer System (Bio-Rad) as per the manufacturer’s instructions. Membranes were immunoblotted as described below.

### Immunoblotting and detection of BN-PAGE and SDS–PAGE assays

Blotted membranes were blocked with 5% milk in Tris-buffered saline plus 0.01% Tween 20 (TBS-T) for an hour at room temperature and subsequently incubated with desired antibodies at 4°C overnight. Antibodies used were anti-GFP (B-2) HRP (Santa Cruz Biotechnology), anti-HA (3F10) HRP (Roche), anti-Myc (9E10) HRP (Roche), anti-FLAG (M2) HRP (Sigma), and anti-V5 (V2260) HRP (Roche), all used in a 1:5,000 dilution in 5% milk in TBS-T. To visualize proteins, we used Pierce ECL Western (32106, Thermo Fisher Scientific), supplementing with up to 50% SuperSignal West Femto Maximum Sensitivity Substrate (34095, Thermo Fishes Scientific) when necessary. Membrane imaging was carried out with an ImageQuant LAS 4000 or an ImageQuant 800 luminescent imager (GE Healthcare Life Sciences, Piscataway, NJ). Rubisco loading control was stained using Ponceau S (Sigma) or Ponceau 4R (Irn Bru, AG Barr).

### Data availability

All relevant data are within the article and in the Supplementary materials. This study includes no data that would need to be deposited in external repositories.

## Acknowledgements

We thank D. Lüdke and C. Marchal for valuable comments on this article. We thank members of the Jones lab (The Sainsbury Laboratory), including Camille-Madeleine Szymansky, Hee-Kyung Ahn and Jonathan Jones, for valuable scientific discussions. We thank Tania Toruño and Peter-Paul Damen (Rijk Zwaan, Netherlands) for providing seeds of lettuce (*L. sativa*) var. ‘Fenston’ and valuable scientific discussions. We thank all members of the TSL Support Services for their invaluable assistance. M.P.C. thanks S. Scorza for support and L. A. “El Flaco” Spinetta for inspiration. The Kamoun Lab is funded primarily from the Gatsby Charitable Foundation, Biotechnology and Biological Sciences Research Council (BBSRC, UK, BB/WW002221/1, BB/V002937/1, BBS/E/J/000PR9795 and BBS/E/J/000PR9796) and the European Research Council (BLASTOFF).

## Author contributions

**Mauricio P Contreras:** Conceptualization; data curation; formal analysis; supervision; validation; investigation; visualization; methodology; writing – original draft; project administration; writing— review and editing. **Hsuan Pai:** Conceptualization; Data curation; formal analysis; supervision; validation; investigation; methodology; writing—review and editing. **Rebecca Thompson:** Data curation; formal analysis; validation; investigation; visualization. **Jules Claeys:** Data curation; formal analysis; validation; investigation. **Hiroaki Adachi:** Conceptualization; investigation; writing—review and editing. **Sophien Kamoun:** Conceptualization; resources; supervision; funding acquisition; visualization; writing—original draft; project administration; writing—review and editing.

## Supplementary Figures

**Figure S1:**
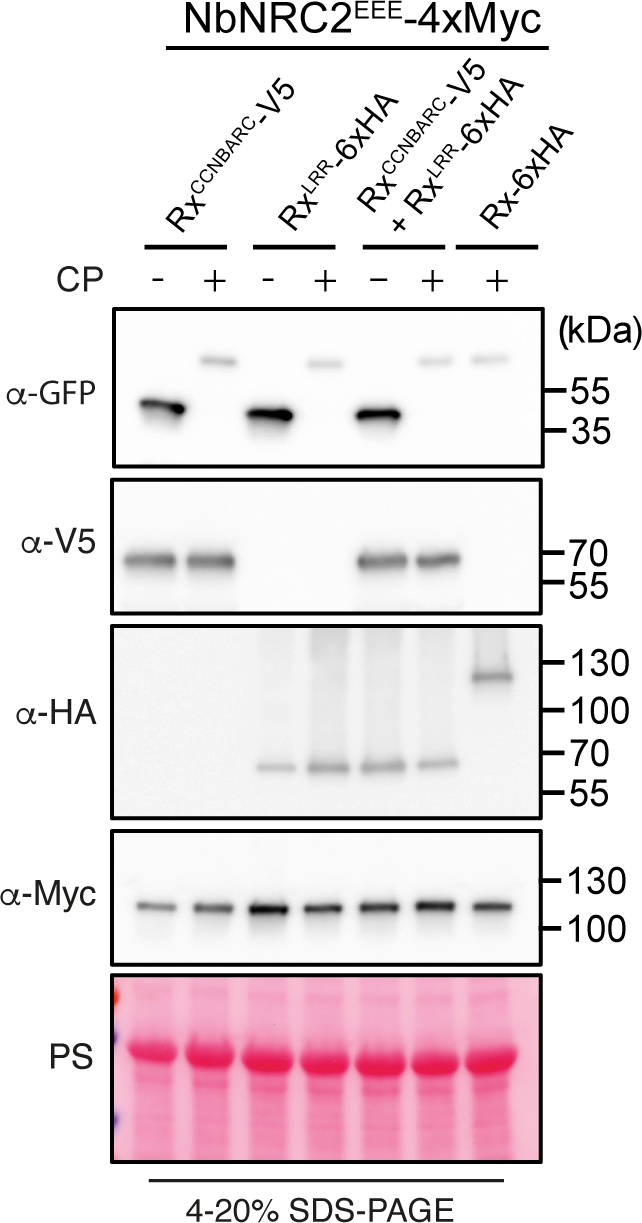
SDS-PAGE blots accompanying BN-PAGE assays in Figure 2. Protein extracts were run on SDS-PAGE assays and immunoblotted with the appropriate antisera labelled on the left. Approximate molecular weights (kDa) of the proteins are shown on the right. Rubisco loading control was carried out using Ponceau stain (PS). Experiment was repeated three times with similar results.

**Figure S2:**
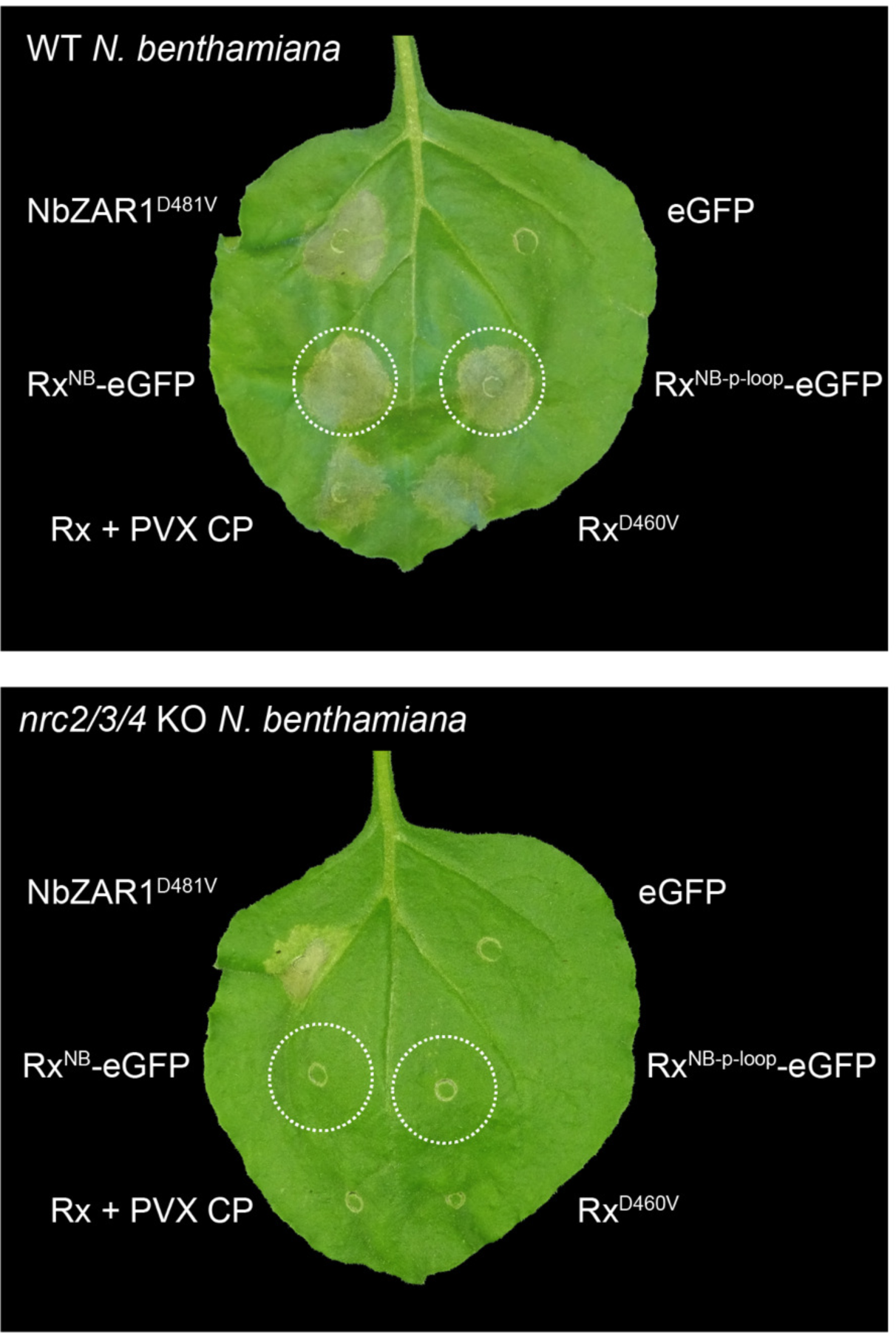
Cell death triggered by Rx^NB^-eGFP does not require an intact p-loop. Photo of representative leaves of WT and *nrc2/3/4* KO *N. benthamiana* plants expressing Rx_NB_-eGFP and an Rx_NB_-eGFP variant with a mutation in the conserved p-loop motif (K to R amino acid substitution). Experiment was repeated three times with at least 6 technical replicates for each repeat. All replicates showed similar results.

**Figure S3:**
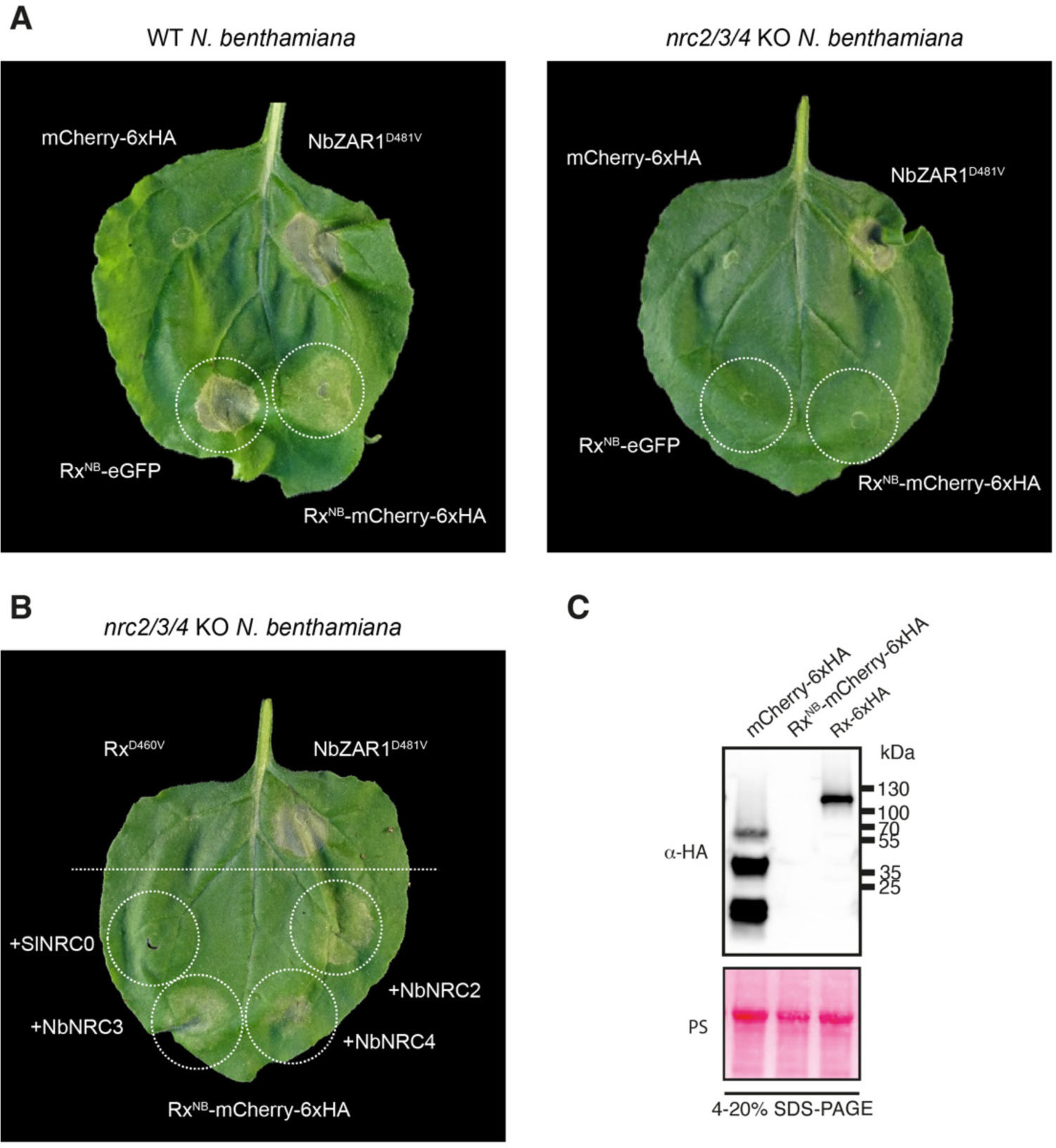
Rx^NB^-mCherry-6xHA also triggers NRC2/3/4-dependent cell death. (**A**) Photo of representative leaves of WT and *nrc2/3/4* KO *N. benthamiana* plants expressing Rx_NB_-mCherry-6xHA. NbZAR1_D481V_ and Rx_NB_-eGFP were included as controls for NRC-independent and NRC-dependent cell death, respectively. mCherry-6xHA was included as a negative control for cell death. Images were taken 5-7 days post agroinfiltration. (**B**) Photo of representative leaves of *nrc2/3/4* KO *N. benthamiana* plants co-expressing Rx_NB_-mCherry-6xHA with NbNRC2, NbNRC3 or NbNRC4. Complementation with SlNRC0 was included as a negative control. NbZAR1_D481V_ and Rx_D460V_ were included as controls for NRC-independent and NRC-dependent cell death, respectively. Images were taken 5 days post agroinfiltration. (**C**) Protein extracts were run on SDS-PAGE assays and immunoblotted with the appropriate antisera labelled on the left. Approximate molecular weights (kDa) of the proteins are shown on the right. Rubisco loading control was carried out using Ponceau stain (PS). Experiment was repeated three times with similar results.

**Figure S4:**
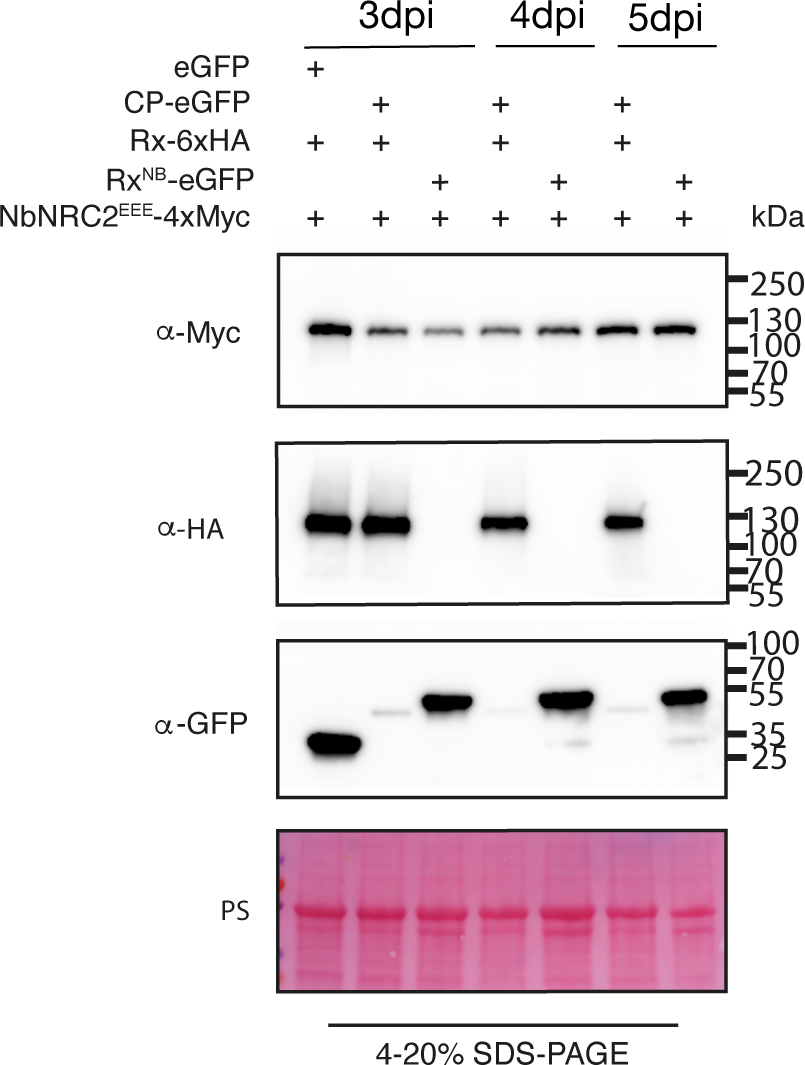
SDS-PAGE blots accompanying BN-PAGE assays in Figure 3. Protein extracts were run on SDS-PAGE assays and immunoblotted with the appropriate antisera labelled on the left. Approximate molecular weights (kDa) of the proteins are shown on the right. Rubisco loading control was carried out using Ponceau stain (PS). Experiment was repeated three times with similar results.

**Figure S5:**
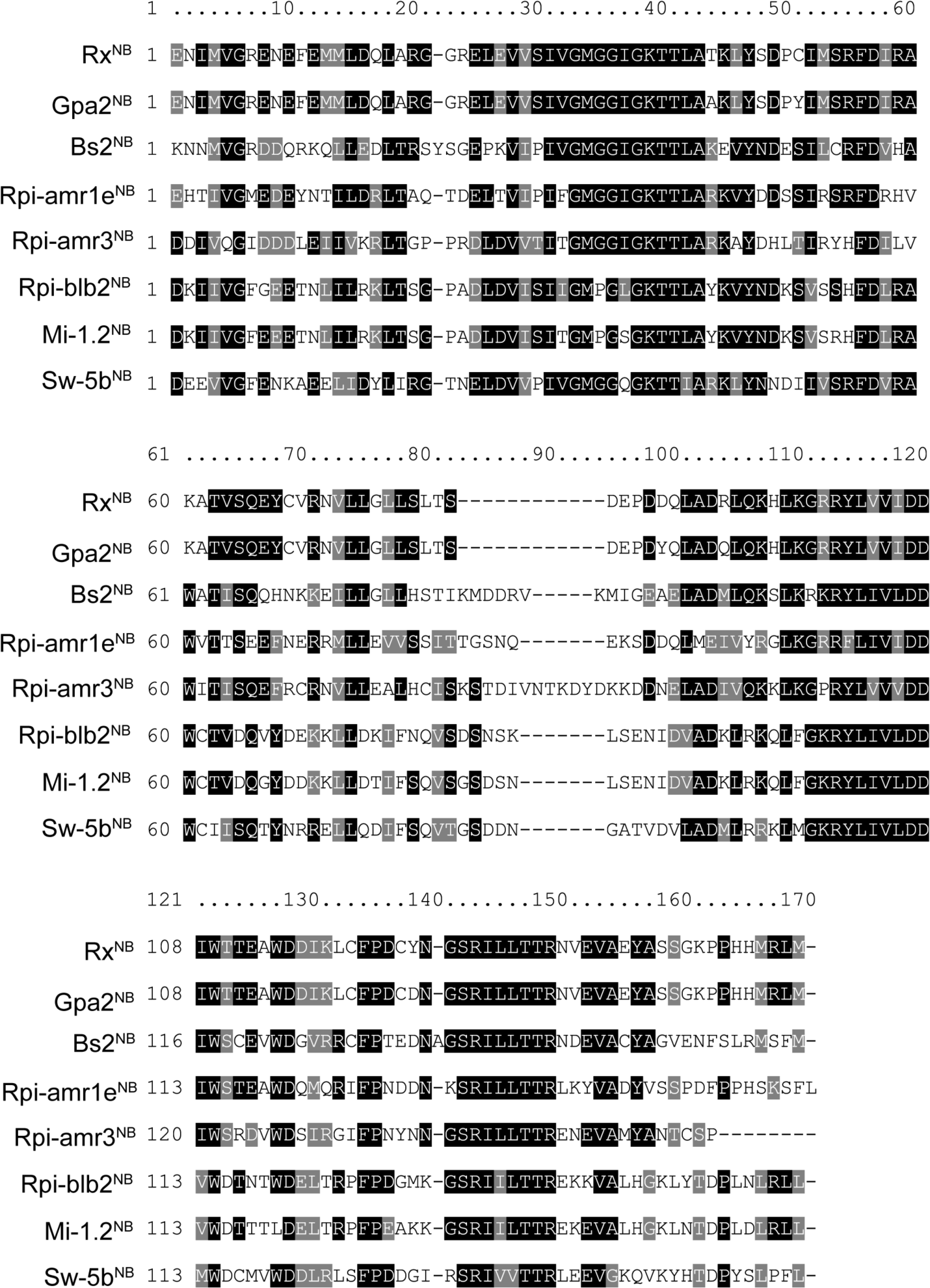
Amino acid sequence alignment of the NB domains of the sensor NLRs Rx, Gpa2, Bs2, Rpi-amr1e, Rpi-amr3, Rpi-blb2, Mi-1.2 and Sw-5b. NB domain boundaries of sensors cloned in this study were determined via alignment to the original Rx_NB_ domain truncation reported by Rairdan and colleagues (Rairdan *et al*., 2008). Amino acid positions indicated correspond to the NB domain alone, with residue in position 1 in Rx_NB_ being equivalent to residue in position 139 for full-length Rx. Alignments were generated using Clustal Omega (Sievers *et al*, 2011).

**Figure S6:**
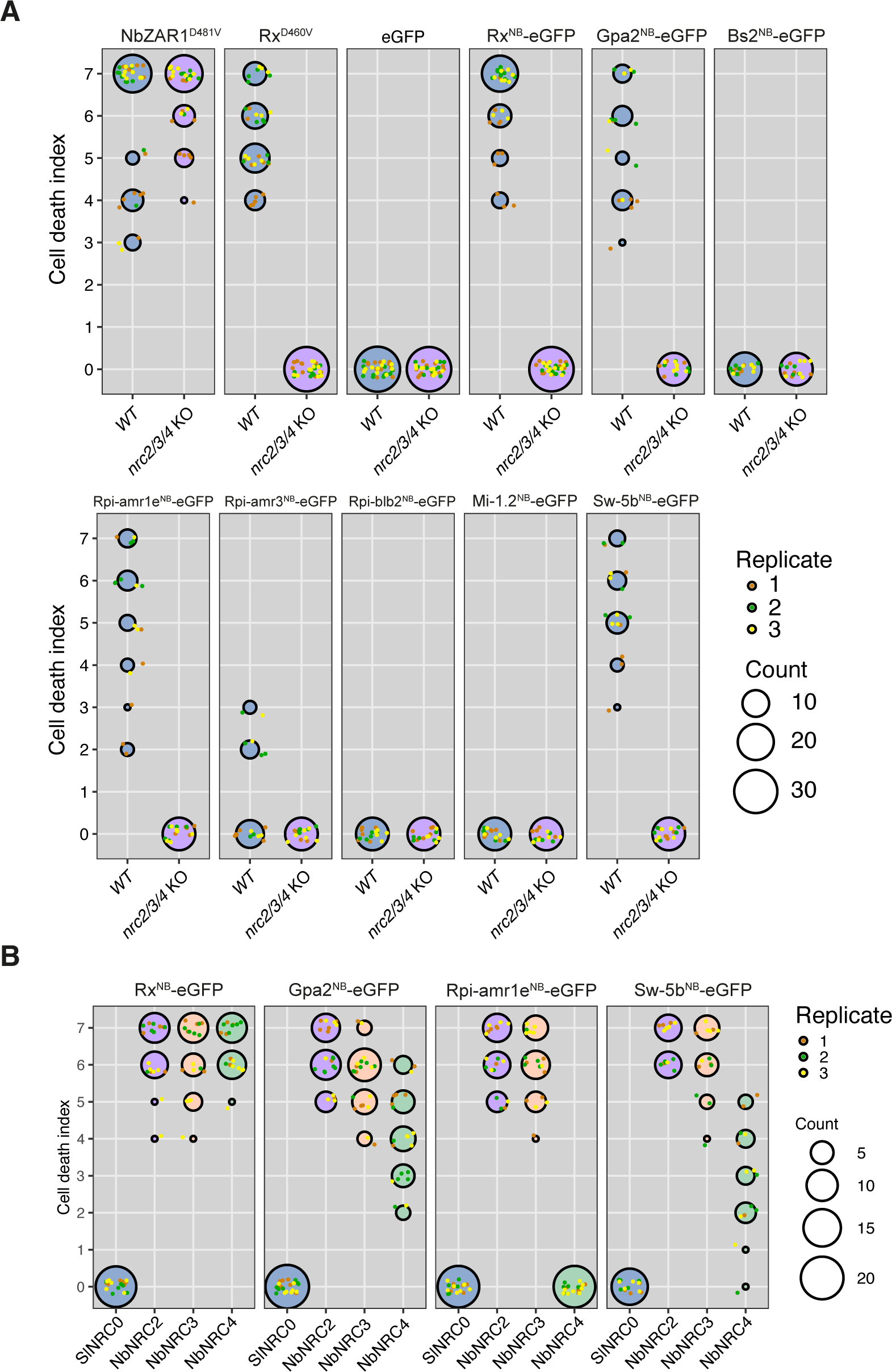
The NB domains of the sensor NLRs Gpa2, Rpi-amr1e, Rpi-amr3 and Sw-5b are sufficient to activate downstream NRC helpers. (**A**) Quantitative analysis of cell death assays shown in **Figure 4B**. (**B**) Quantitative analysis of cell death assays shown in **Figure 3C and Figure 4C**. (**A-B**) HR cell death was scored based on a modified 0-7 scale (Segretin *et al*, 2014) at 5-7 days post agroinfiltration. HR scores are presented as dot plots, where the size of each dot is proportional to the number of samples with the same score (Count). Results are based on 3 biological replicates.

**Figure S7:**
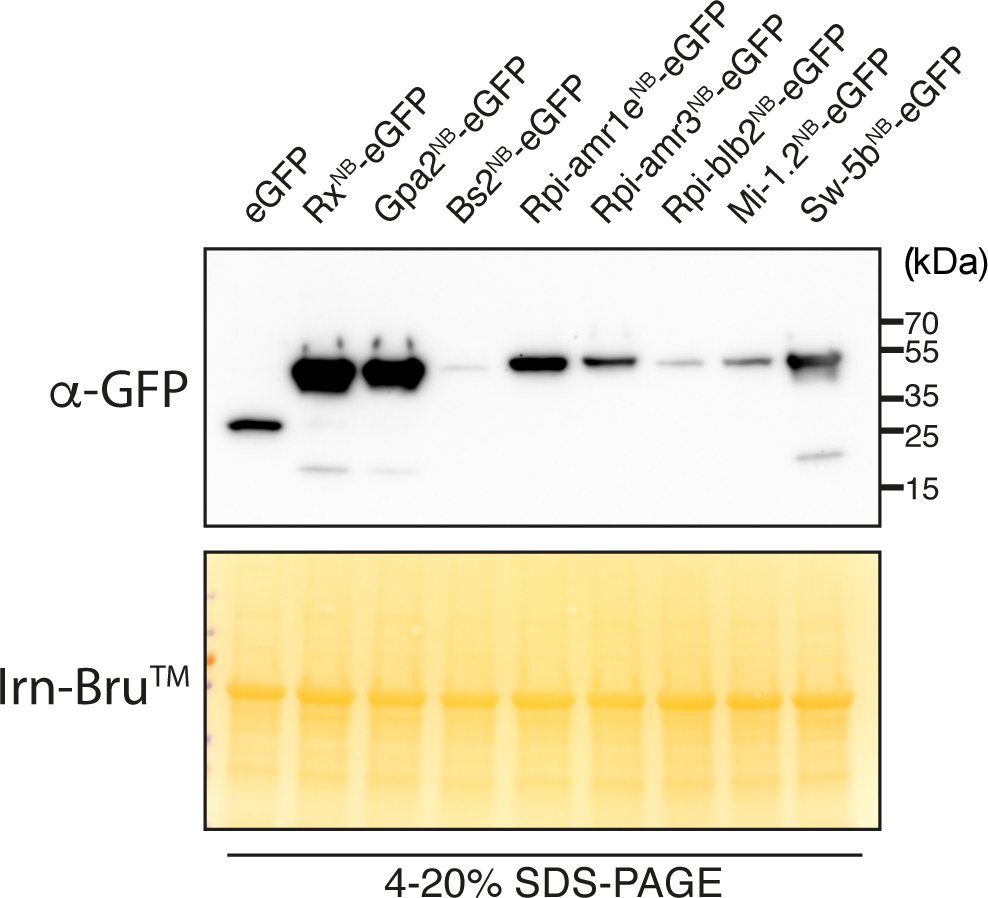
All cell death-inducing sensor NLR NB domain-eGFP fusions accumulate in planta. Protein extracts from leaves of *nrc2/3/4* KO *N. benthamiana* expressing proteins of interest were run on SDS-PAGE assays and immunoblotted with the appropriate antisera labelled on the left. Approximate molecular weights (kDa) of the proteins are shown on the right. Rubisco loading control was carried out using Ponceau 4R staining (Irn-Bru_TM_). Experiment was repeated two times with similar results.

**Figure S8:**
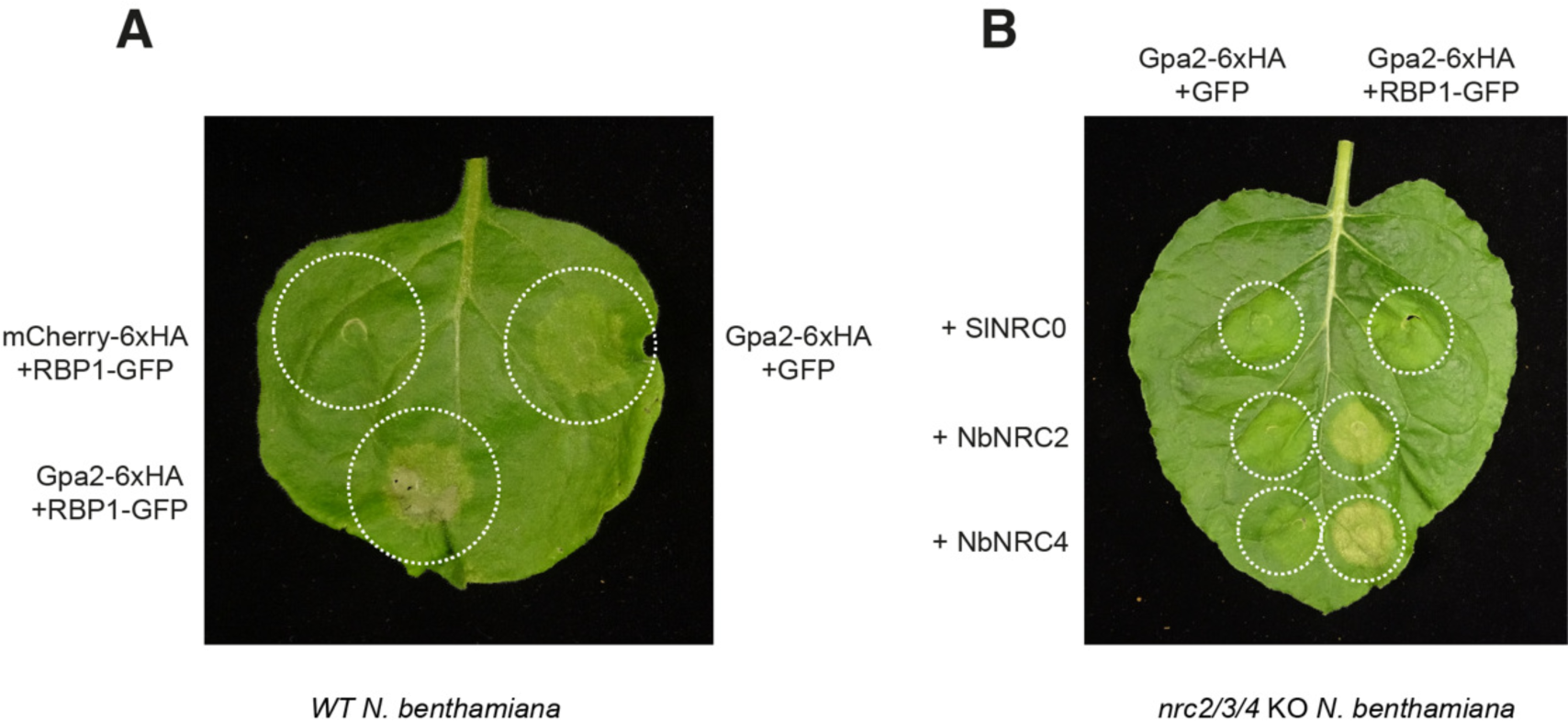
The full-length sensor NLR Gpa2 can signal through NRC2 and NRC4. (**A**) Photo of a representative leaf of WT *N. benthamiana* showing cell death after co-expression of Gpa2 and RBP1. (**B**) Photo of a representative leaf of *nrc2/3/4* KO *N. benthamiana* showing cell death after co-expression of Gpa2/RBP1 and NRC2 or NRC4. The *S. lycopersicum* helper SlNRC0 was included as a negative control for complementation. GFP was used as a negative control for RBP1-GFP. (**A-B**) Images were taken 5-7 days after agroinfiltration. Experiment was repeated 2 times with similar results.

## References

Adachi H, Contreras MP, Harant A, Wu C-h, Derevnina L, Sakai T, Duggan C, Moratto E, Bozkurt TO, Maqbool A et al (2019a) An N-terminal motif in NLR immune receptors is functionally conserved across distantly related plant species. Elife 8: e49956

Adachi H, Derevnina L, Kamoun S (2019b) NLR singletons, pairs, and networks: evolution, assembly, and regulation of the intracellular immunoreceptor circuitry of plants. Current opinion in plant biology 50: 121–131

Adachi H, Kamoun S (2022) NLR receptor networks in plants. Essays in Biochemistry 66: 541–549

Ahn HK, Lin X, Olave-Achury AC, Derevnina L, Contreras MP, Kourelis J, Wu CH, Kamoun S, Jones JD (2023) Effector-dependent activation and oligomerization of plant NRC class helper NLRs by sensor NLR immune receptors Rpi-amr3 and Rpi-amr1. The EMBO Journal: e111484

Bendahmane A, Kanyuka K, Baulcombe DC (1999) The Rx gene from potato controls separate virus resistance and cell death responses. The Plant Cell 11: 781–791

Bendahmane A, Köhm BA, Dedi C, Baulcombe DC (1995) The coat protein of potato virus X is a strain-specific elicitor of Rx1-mediated virus resistance in potato. The Plant Journal 8: 933–941

Bentham AR, Zdrzałek R, De la Concepcion JC, Banfield MJ (2018) Uncoiling CNLs: structure/function approaches to understanding CC domain function in plant NLRs. Plant and Cell Physiology 59: 2398–2408

Bi G, Su M, Li N, Liang Y, Dang S, Xu J, Hu M, Wang J, Zou M, Deng Y (2021) The ZAR1 resistosome is a calcium-permeable channel triggering plant immune signaling. Cell 184: 3528–3541. e3512

Chou W-C, Jha S, Linhoff MW, Ting JP-Y (2023) The NLR gene family: from discovery to present day. Nature Reviews Immunology: 1–20

Contreras MP, Kamoun S (2022) The high molecular weight complex formed by the plant immune receptor Rx does not contain host protein RanGAP2. Zenodo

Contreras MP, Lüdke D, Pai H, Toghani A, Kamoun S (2023a) NLR receptors in plant immunity: making sense of the alphabet soup. EMBO reports: e57495

Contreras MP, Pai H, Selvaraj M, Toghani A, Lawson DM, Tumtas Y, Duggan C, Yuen ELH, Stevenson CEM, Harant A et al (2023b) Resurrection of plant disease resistance proteins via helper NLR bioengineering. Science Advances 9: eadg3861

Contreras MP, Pai H, Tumtas Y, Duggan C, Yuen ELH, Cruces AV, Kourelis J, Ahn HK, Lee KT, Wu CH et al (2023c) Sensor NLR immune proteins activate oligomerization of their NRC helpers in response to plant pathogens. The EMBO Journal 42: e111519

Dangl JL, Jones JD (2001) Plant pathogens and integrated defence responses to infection. nature 411: 826–833

Derevnina L, Contreras MP, Adachi H, Upson J, Vergara Cruces A, Xie R, Skłenar J, Menke FL, Mugford ST, MacLean D et al (2021) Plant pathogens convergently evolved to counteract redundant nodes of an NLR immune receptor network. PLoS biology 19: e3001136

Duxbury Z, Wu C-h, Ding P (2021) A comparative overview of the intracellular guardians of plants and animals: NLRs in innate immunity and beyond. Annual review of plant biology 72: 155–184

Engler C, Youles M, Gruetzner R, Ehnert T-M, Werner S, Jones JD, Patron NJ, Marillonnet S (2014) A golden gate modular cloning toolbox for plants. ACS synthetic biology 3: 839–843

Förderer A, Kourelis J (2023) NLR immune receptors: structure and function in plant disease resistance. Biochemical Society Transactions 51: 1473–1483

Förderer A, Li E, Lawson AW, Deng Y-n, Sun Y, Logemann E, Zhang X, Wen J, Han Z, Chang J et al (2022) A wheat resistosome defines common principles of immune receptor channels. Nature 610: 532–539

Goh F-J, Huang C-Y, Derevnina L, Wu C-H (2023) NRC immune receptor networks show diversified hierarchical genetic architecture across plant lineages. bioRxiv: 2023.2010. 2025.563953

Harant A, Pai H, Sakai T, Kamoun S, Adachi H (2022) A vector system for fast-forward studies of the HOPZ-ACTIVATED RESISTANCE1 (ZAR1) resistosome in the model plant Nicotiana benthamiana. Plant Physiology 188: 70–80

Huang S, Jia A, Song W, Hessler G, Meng Y, Sun Y, Xu L, Laessle H, Jirschitzka J, Ma S (2022) Identification and receptor mechanism of TIR-catalyzed small molecules in plant immunity. Science 377: eabq3297

Kourelis J, Contreras MP, Harant A, Pai H, Lüdke D, Adachi H, Derevnina L, Wu C-H, Kamoun S (2022) The helper NLR immune protein NRC3 mediates the hypersensitive cell death caused by the cell-surface receptor Cf-4. PLoS Genetics 18: e1010414

Kourelis J, Sakai T, Adachi H, Kamoun S (2021) RefPlantNLR is a comprehensive collection of experimentally validated plant disease resistance proteins from the NLR family. PLoS Biology 19: e3001124

Lin X, Olave-Achury A, Heal R, Pais M, Witek K, Ahn H-K, Zhao H, Bhanvadia S, Karki HS, Song T et al (2022) A potato late blight resistance gene protects against multiple Phytophthora species by recognizing a broadly conserved RXLR-WY effector. Molecular Plant 15: 1457–1469

Ma S, Lapin D, Liu L, Sun Y, Song W, Zhang X, Logemann E, Yu D, Wang J, Jirschitzka J et al (2020) Direct pathogen-induced assembly of an NLR immune receptor complex to form a holoenzyme. Science 370: eabe3069

Marchal C, Pai H, Kamoun S, Kourelis J (2022) Emerging principles in the design of bioengineered made-to-order plant immune receptors. Current Opinion in Plant Biology: 102311

Mestre P, Baulcombe DC (2006) Elicitor-mediated oligomerization of the tobacco N disease resistance protein. The Plant Cell 18: 491–501

Meyers BC, Shen KA, Rohani P, Gaut BS, Michelmore RW (1998) Receptor-like genes in the major resistance locus of lettuce are subject to divergent selection. The Plant Cell 10: 1833–1846

Moffett P, Farnham G, Peart J, Baulcombe DC (2002) Interaction between domains of a plant NBS–LRR protein in disease resistance-related cell death. The EMBO journal 21: 4511–4519

Rairdan GJ, Collier SM, Sacco MA, Baldwin TT, Boettrich T, Moffett P (2008) The coiled-coil and nucleotide binding domains of the potato Rx disease resistance protein function in pathogen recognition and signaling. The Plant Cell 20: 739–751

Sakai T, Martinez-Anaya C, Contreras MP, Kamoun S, Wu C-H, Adachi H (2023) The NRC0 gene cluster of sensor and helper NLR immune receptors is functionally conserved across asterid plants. bioRxiv: 2023.2010. 2023.563533

Segretin ME, Pais M, Franceschetti M, Chaparro-Garcia A, Bos JI, Banfield MJ, Kamoun S (2014) Single amino acid mutations in the potato immune receptor R3a expand response to Phytophthora effectors. Molecular Plant-Microbe Interactions 27: 624–637

Shen KA, Chin DB, Arroyo-Garcia R, Ochoa OE, Lavelle DO, Wroblewski T, Meyers BC, Michelmore RW (2002) Dm3 is one member of a large constitutively expressed family of nucleotide binding site— leucine-rich repeat encoding genes. Molecular plant-microbe interactions 15: 251–261

Sievers F, Wilm A, Dineen D, Gibson TJ, Karplus K, Li W, Lopez R, McWilliam H, Remmert M, Söding J (2011) Fast, scalable generation of high-quality protein multiple sequence alignments using Clustal Omega. Molecular systems biology 7: 539

Tameling WI, Baulcombe DC (2007) Physical association of the NB-LRR resistance protein Rx with a Ran GTPase–activating protein is required for extreme resistance to Potato virus X. The Plant Cell 19: 1682–1694

Van Der Biezen EA, Jones JD (1998) Plant disease-resistance proteins and the gene-for-gene concept. Trends in biochemical sciences 23: 454–456

van der Hoorn RA, Kamoun S (2008) From guard to decoy: a new model for perception of plant pathogen effectors. The Plant Cell 20: 2009–2017

Wang J, Hu M, Wang J, Qi J, Han Z, Wang G, Qi Y, Wang H-W, Zhou J-M, Chai J (2019) Reconstitution and structure of a plant NLR resistosome conferring immunity. Science 364: eaav5870

Weber E, Engler C, Gruetzner R, Werner S, Marillonnet S (2011) A modular cloning system for standardized assembly of multigene constructs. PloS one 6: e16765

Witek K, Lin X, Karki HS, Jupe F, Witek AI, Steuernagel B, Stam R, Van Oosterhout C, Fairhead S, Heal R (2021) A complex resistance locus in Solanum americanum recognizes a conserved Phytophthora effector. Nature Plants 7: 198–208

Wróblewski T, Spiridon L, Martin EC, Petrescu A-J, Cavanaugh K, Truco MJ, Xu H, Gozdowski D, Pawłowski K, Michelmore RW (2018) Genome-wide functional analyses of plant coiled–coil NLR-type pathogen receptors reveal essential roles of their N-terminal domain in oligomerization, networking, and immunity. PLoS biology 16: e2005821

Wu C-H, Abd-El-Haliem A, Bozkurt TO, Belhaj K, Terauchi R, Vossen JH, Kamoun S (2017) NLR network mediates immunity to diverse plant pathogens. Proceedings of the National Academy of Sciences 114: 8113–8118

Wu C-H, Derevnina L, Kamoun S (2018) Receptor networks underpin plant immunity. Science 360: 1300–1301

Zeng L, Zhang Q, Sun R, Kong H, Zhang N, Ma H (2014) Resolution of deep angiosperm phylogeny using conserved nuclear genes and estimates of early divergence times. Nature communications 5: 4956

Zhang L, Chen S, Ruan J, Wu J, Tong AB, Yin Q, Li Y, David L, Lu A, Wang WL (2015) Cryo-EM structure of the activated NAIP2-NLRC4 inflammasome reveals nucleated polymerization. Science 350: 404–409

Zhao Y-B, Liu M-X, Chen T-T, Ma X, Li Z-K, Zheng Z, Zheng S-R, Chen L, Li Y-Z, Tang L-R (2022) Pathogen effector AvrSr35 triggers Sr35 resistosome assembly via a direct recognition mechanism. Science Advances 8: eabq5108

